# Genetic parallels in biomineralization of the calcareous sponge *Sycon ciliatum* and stony corals

**DOI:** 10.1101/2025.02.06.636789

**Authors:** Oliver Voigt, Magdalena V. Wilde, Thomas Fröhlich, Benedetta Fradusco, Sergio Vargas, Gert Wörheide

**Affiliations:** Department of Earth and Environmental Sciences, Paleontology and Geobiology, Ludwig-Maximilians-Universität München, Munich, Germany; Gene Center - Laboratory for Functional Genome Analysis, Ludwig-Maximilians-Universität München, Munich, Germany; GeoBio-Center, Ludwig-Maximilians-Universität München, Munich, Germany; Staatliche Naturwissenschaftliche Sammlungen Bayerns (SNSB)–Bayerische Staatssammlung für Paläontologie und Geologie, Munich, Germany

**Keywords:** Biomineralization, gene duplication, convergent evolution, calcareous sponges, corals

## Abstract

The rapid emergence of mineralized structures in diverse animal groups during the late Ediacaran and early Cambrian periods likely resulted from modifications of pre-adapted biomineralization genes inherited from a common ancestor. As the oldest extant phylum with mineralized structures, sponges are key to understanding animal biomineralization. Yet, the biomineralization process in sponges, particularly in forming spicules, is not well-understood. To address this, we conducted transcriptomic, genomic, and proteomic analyses on the calcareous sponge *Sycon ciliatum*, supplemented by in situ hybridization. We identified 829 genes overexpressed in regions of increased calcite spicule formation, including 17 calcarins—proteins analogous to corals’ galaxins localized in the spicule matrix and expressed in sclerocytes. Their expression varied temporally and spatially, specific to certain spicule types, indicating that fine-tuned gene regulation is crucial for biomineralization control. Similar subtle expression changes are also relevant in stony coral biomineralization. Tandem gene arrangements and expression changes suggest that gene duplication and neofunctionalization have significantly shaped *Sycon ciliatum*’s biomineralization, similar to that in corals. These findings suggest a parallel evolution of carbonate biomineralization in the calcitic *Sycon ciliatum* and aragonitic corals, exemplifying the evolution of mechanisms crucial for animals to act as ecosystem engineers and form reef structures.

## Introduction

The evolution of biomineralization allowed animals to produce important mineralized functional structures, such as shells, teeth, and skeletons. For this, they most frequently use calcium carbonate, calcium phosphate, and silicate as mineral components, of which calcium carbonate is taxonomically the most diverse (Knoll, 2003; Murdock and Donoghue, 2011). Animal biominerals are usually composite materials of organic and mineral compounds and exhibit material properties and shapes that differ enormously from their purely mineral counterparts. Their formation requires biological control governed by specific genes and proteins in specialized biomineralization cells and tissues. Those then ensure supply with the necessary inorganic substances into the calcifying space (intracellularly or extracellularly) to enable crystal precipitation and growth, e.g., by pH regulation. They also provide organic components, including secreted, so-called “matrix proteins” that influence the polymorph and shape of the biomineral and subsequently may become embedded in it (Aizenberg et al., 1996, 1994).

The ability to form biominerals evolved several times independently in animal lineages, and early instances of mineralized animal skeletons appeared within a geologically brief span, from the latest Ediacaran to the Middle Cambrian (Murdock and Donoghue, 2011). According to the ‘biomineralization toolkit’ hypothesis, animals of this era had acquired pre-adapted genes and gene-regulatory networks necessary to produce biominerals in a controlled manner that were further refined independently in different lineages (Murdock, 2020). This can be achieved by gene family expansion, in which, after initial gene duplication, diversification by mutation and natural selection leads to the neofunctionalization of a copied gene in the biomineralization process.

Among early branching, non-bilaterian calcifiers, stony corals are best studied, and essential elements of their genetic biomineralization machinery are known that originate from gene duplication and neofunctionalization, such as the SLC4ɣ bicarbonate transporter (SLC4ɣ) that is a key component of the coral aragonite skeleton forming machinery (Tinoco et al., 2023; Zoccola et al., 2015). Numerous skeletal matrix proteins have been identified (Drake et al., 2013; Peled et al., 2020; Ramos-Silva et al., 2013). Among those, galaxin and related proteins were the first to be characterized, and at least in some species, comprise the most dominant protein in skeletal matrix extractions (Fukuda et al., 2003; Watanabe et al., 2003) Acidic proteins are another essential component of coral skeletal matrices, presumably influencing the polymorph of the precipitating carbonate (Mass et al. 2013; Laipnik et al. 2020; Mass et al. 2016). However, the genetic mechanisms of the calcium carbonate biomineralization in sponges (class Porifera) are less known. Sponges produce skeletal elements in the form of differently shaped spicules, siliceous in the extant sponge classes Demospongiae, Hexactinellida, and Homoscleromorpha, and calcitic only in the class Calcarea (calcareous sponges). Nonetheless, some demosponges like the polyphyletic “sclerosponges,” may form a rigid calcium carbonate basal skeleton in addition to or instead of siliceous spicules (Wörheide, 1998). Essential skeletal matrix proteins occluded in such rigid calcitic carbonate skeletons of sclerosponges have been studied in *Astrosclera willeyana* (Jackson et al., 2011, 2007) and *Vaceletia* sp.(Germer et al., 2015). In contrast to those aragonite-producing sclerosponges, calcareous sponges continuously produce calcitic spicules of different shapes, providing a unique opportunity to observe all stages of the process and study differences in their production. Additionally, their spicule formation is often faster than the production of skeletal elements in other calcifiers. It takes only a few days from initiation to the finished spicule and involves only a few specialized cells called sclerocytes (Ilan et al., 1996; Woodland, 1905). They control biomineralization by their relative movement to each other and by secreting specific proteins, ions, and other substances into the calcification space, a process still little understood at the molecular level. Only a few genes involved in calcification in calcareous sponges are known, including two specific carbonic anhydrases, and two SLC4 bicarbonate transporters, and three acidic proteins, two of which are spicule-type specific (Voigt et al., 2021, 2017, 2014). Skeletal matrix proteins have not yet been identified but are of particular interest because they may play a crucial role in spicule morphogenesis (Aizenberg et al., 1995).

In this study, we used genomic data, differential gene expression analysis, and RNA *in situ* hybridization experiments, supplemented by a proteomic approach, to identify genes directly involved in the spicule formation in the calcarean model species *Sycon ciliatum* (subclass Calcaronea), and compare those with another non-bilaterian calcifying clade, the scleractinian (stony) corals, the ecosystem engineers of today’s coral reefs. We found surprising similarities in key components of the biomineralization toolkit between calcareous sponges and corals that shed new light on the evolution of calcium carbonate biomineralization in animals.

## Results

*Sycon ciliatum* is a tube-shaped sponge with a single apical osculum and a sponge wall of radial tubes around the central atrium (Fig. 1A). The radial tubes are internally lined with choanoderm, which forms elongated chambers in an angle of approximately 90° to the tube axis. The sponge tissue is supported by a skeleton composed of countless spicules arranged in a specific pattern within the sponge body (Fig. 1 A). The spicules are formed by sclerocytes located within the mesohyl, the extracellular matrix of the sponge. Connected by septate junctions, they enclose an extracellular space where a spicule grows (Ledger, 1975). Only a small number of sclerocytes participate in forming a single spicule (Fig. 1C). Production of each actine (ray) of a spicule involves a pair of sclerocytes (Woodland, 1905): a founder cell, which initiates actine elongation at the tip, and a thickener cell, located alongside the actine. The thickener cell enhances actine strength in some species by precipitating additional calcium carbonate. More spicules are produced in the apical growth zone than in other regions of *Sycon’s* body (Voigt et al., 2014). In this zone, diactine spicules, one-rayed spicules with two pointed tips, are arranged in a palisade-like manner and continuously grow around the osculum opening. Simultaneously, new triactines, three-rayed spicules resembling a Mercedes star, and four-rayed tetractines form in the atrial skeleton. Large amounts of triactines and distal diactine bundles are also produced within newly developing radial tubes (Fig.1B).

**Figure 1.**
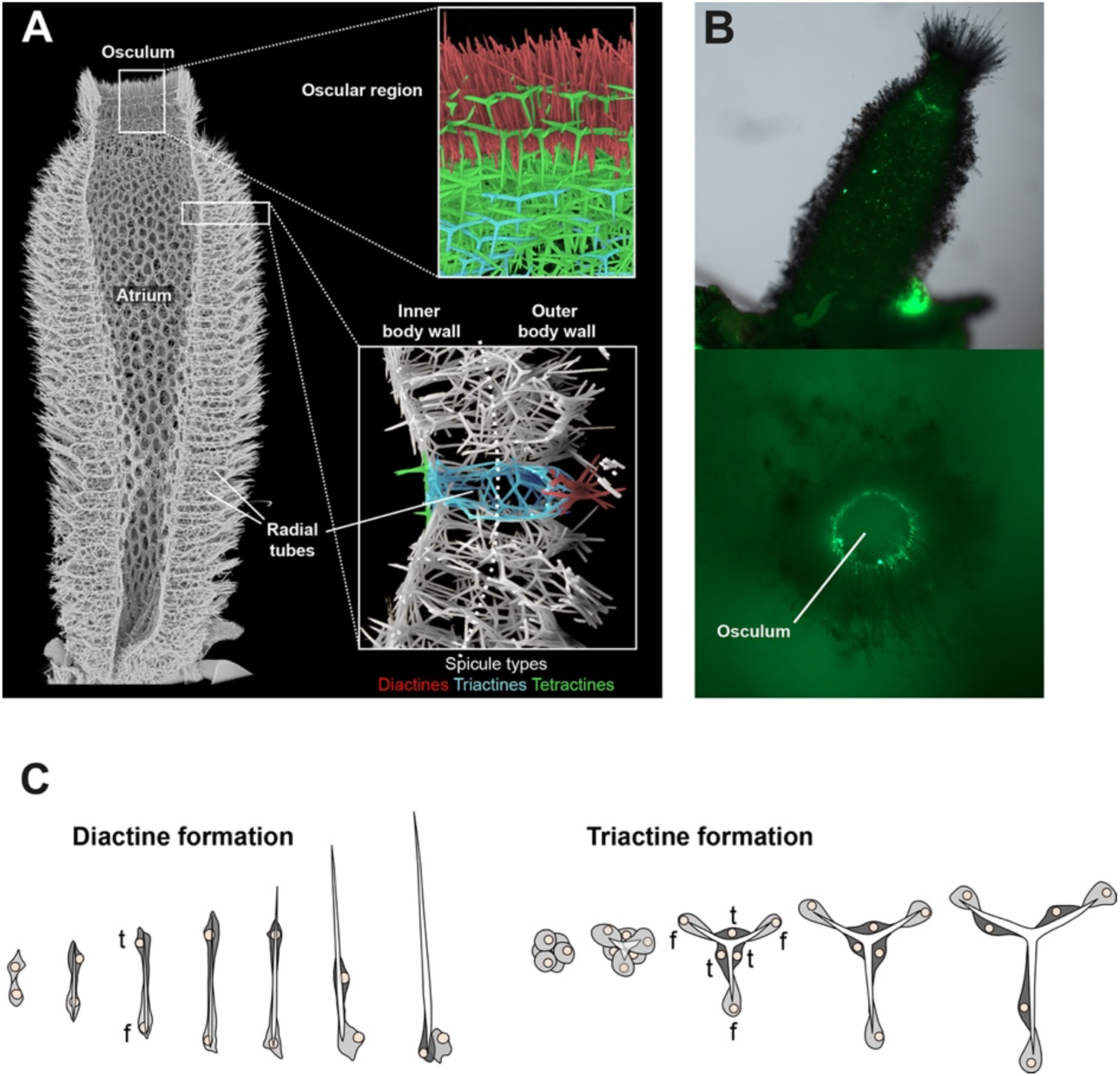
The *Sycon ciliatum* skeleton features specific spicule types in distinct body regions: parallel diactines in the oscular region (upper inset), radial tubes supported by triactines and tufted with diactines (lower inset), and the atrial skeleton composed of triactines and tetractines. B: The upper oscular region shows increased spicule formation (calcein staining) in the growing zone of new radial tubes and around the osculum, where oscular diactines are predominantly produced (modified from Voigt et al., 2014). C: Spicules are formed by sclerocytes, specialized cells controlling spicule formation. Diactine formation involves two sclerocytes, triactine formation six (f = founder cell, t = thickener cell).

### Differential gene expression analysis

We performed a differential gene expression analysis comparing gene expression of the apical oscular region to the inner and outer body walls of the more basal body regions in five specimens (Table S1). We observed 1,575 differentially expressed genes (log2-fold change ≥ 2, padj < 0.01). Of these genes, 829 were over-expressed in the oscular region, including the known biomineralization genes with documented sclerocyte-specific expression: CA1, CA2, AE-like1, AE-like2, Diactinin, Triactinin, Spiculin (Voigt et al., 2017, 2014).

To gain further insight into these genes’ potential function, we performed a GO-term enrichment analysis, considering the GO-term annotation of the closest hit in Uniprot against each transcript as a potential annotation for the *Sycon ciliatum* genes. The genes overexpressed in the oscular regions were enriched in biological process GO-terms relevant for biomineralization (e.g., with the representative terms GO:0001501 “skeletal system development,” GO:0001503: “ossification”, “GO:0030282 “bone mineralization”, Dataset S1). Among them were genes similar to Fibrilin-1, Collagen alpha-1 (XXVIII) chain B, Collagen alpha-1 (I) chain Collagen alpha-1(XI) chain, V-type proton ATPase subunit a, and to Fibroblast growth factor receptor 2 (FGFR2). Additionally, the *Sycon ciliatum* specific growth factor SciTgfBH, with similarity to bone morphogenetic protein 2, and the homeobox protein SciMsx, with similarity to the homeobox protein MSX-2, belong to genes with these enriched GO-terms. Other enriched GO terms related to ongoing morphogenesis, cell differentiation, cell signaling, cell migration, and organization of the organic matrix are all indicative of the growth zone that the oscular region represents (Dataset S2). Noteworthy among them are genes of the Wnt Pathway (SciWntG, I, J, K, L, M, O, U, SciFzdD, and Metalloprotease TIKI homolog).

Fourteen secreted proteins, with significantly higher expression in the oscular region, showed similarity to the coral skeletal organic matrix proteins Galaxin and galaxin-like proteins, first described from the coral *Galaxea fascicularis* (Fukuda et al., 2003; Watanabe et al., 2003). In total, 17 proteins similar to galaxins were identified from the predicted proteins of the *Sycon ciliatum* genome using BLASTp. We refer to these as Calcarin 1 - 17 (Cal1–Cal17) to discriminate them from galaxin and galaxin-like proteins of scleractinian corals. The presence of signal peptides indicates that calcarins are secreted. Except for a Reeler domain (Pfam PF02014) in Cal16, calcarins lack recognizable domains. Like galaxins, calcarins contain regions with 10 to 23 di-cysteine residues, typically separated by 10 to 15 amino acids. Additionally, a single cysteine occurs approximately at the inter-di-cysteine distance upstream and downstream of the di-cysteine region. Monomer Alpha-Fold predictions propose that disulfide bridges between these cysteines provide a tertiary structure typical of calcarins and galaxins (Figs. 2 A, S1). The di-cysteines form the N-terminal part of a common four-amino acid beta-hairpin with a short, often two-amino acid long turn. The first cysteine of a di-cysteine connects with the second cysteine of the preceding beta-hairpin by a disulfide bridge, linking the beta-hairpins together (Fig. 2 B). This disulfide bridge backbone is continuous in galaxins, whereas the predictions in calcarins may show one or two interruptions.

**Figure 2.**
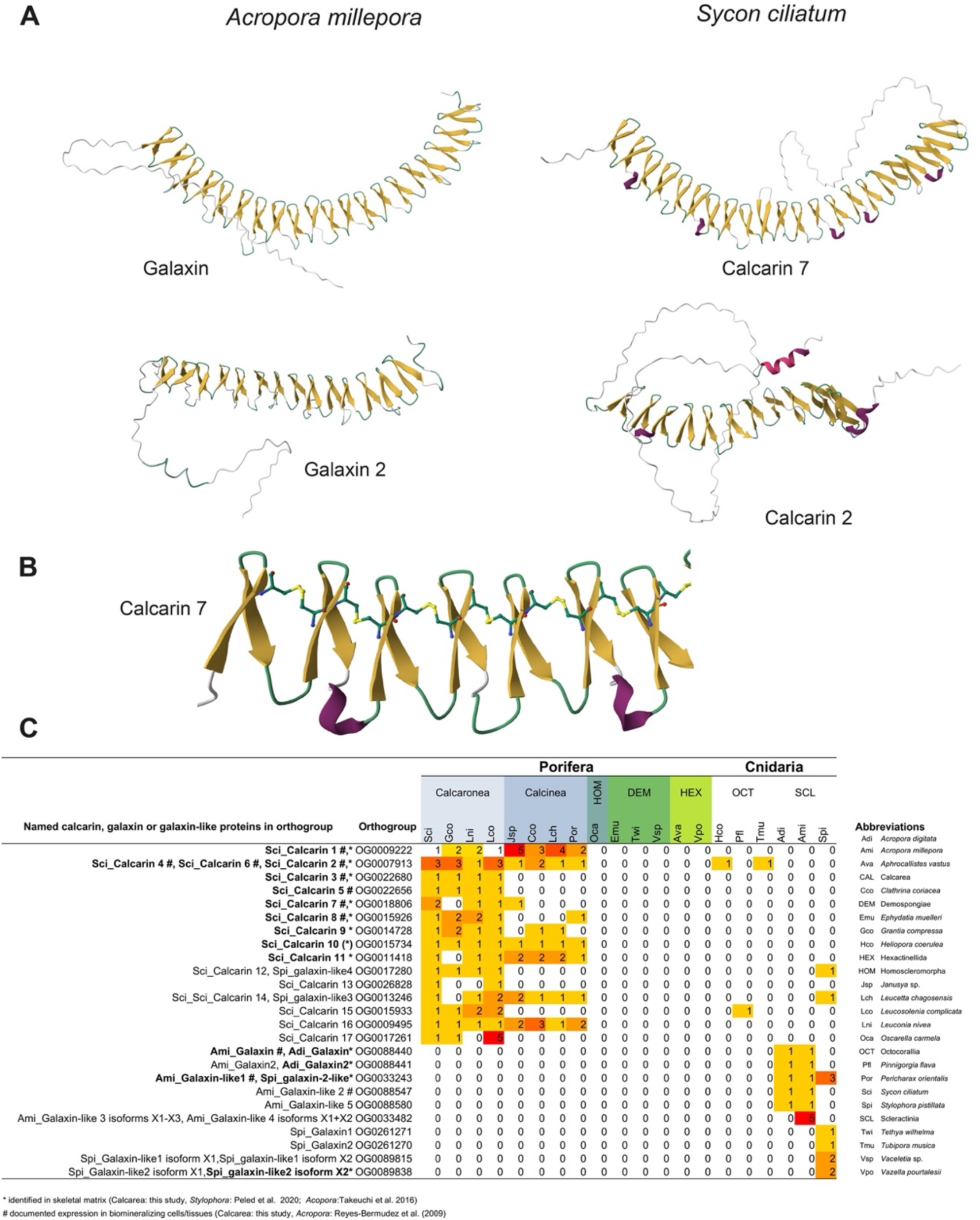
A: Structural similarities in Alpha-Fold predictions of galaxins (*Acropora millepora*) and selected calcarins (*Sycon ciliatum*). B: Beta-hairpins in Cal7 connected by disulfide bridges of di-cysteines. C: Number of calcarins, galaxin-like, and galaxins transcripts in sponges and corals, assigned to orthogroups.

### Temporal and spatial expression of calcarins

The expression of Cal1 to Cal8 was investigated using chromogenic *in situ* hybridization (CISH) and hairpin-chain reaction fluorescence *in situ* hybridization (HCR-FISH), confirming their presence in sclerocytes (Fig. 3). Additionally, antisense probes were used against the mRNA of Spiculin, an acidic protein specific to all thickener cells, Triactinin, a marker for triactine and tetractine thickener cells, and the sclerocyte-specific carbonic anhydrase SciCA1 (Voigt et al., 2017, 2014). This approach enabled us to visualize and contextualize Calcarin-positive cells (Fig. S2). The expression patterns of Calcarins varied with sclerocyte type and spicule formation stage.

**Figure 3.**
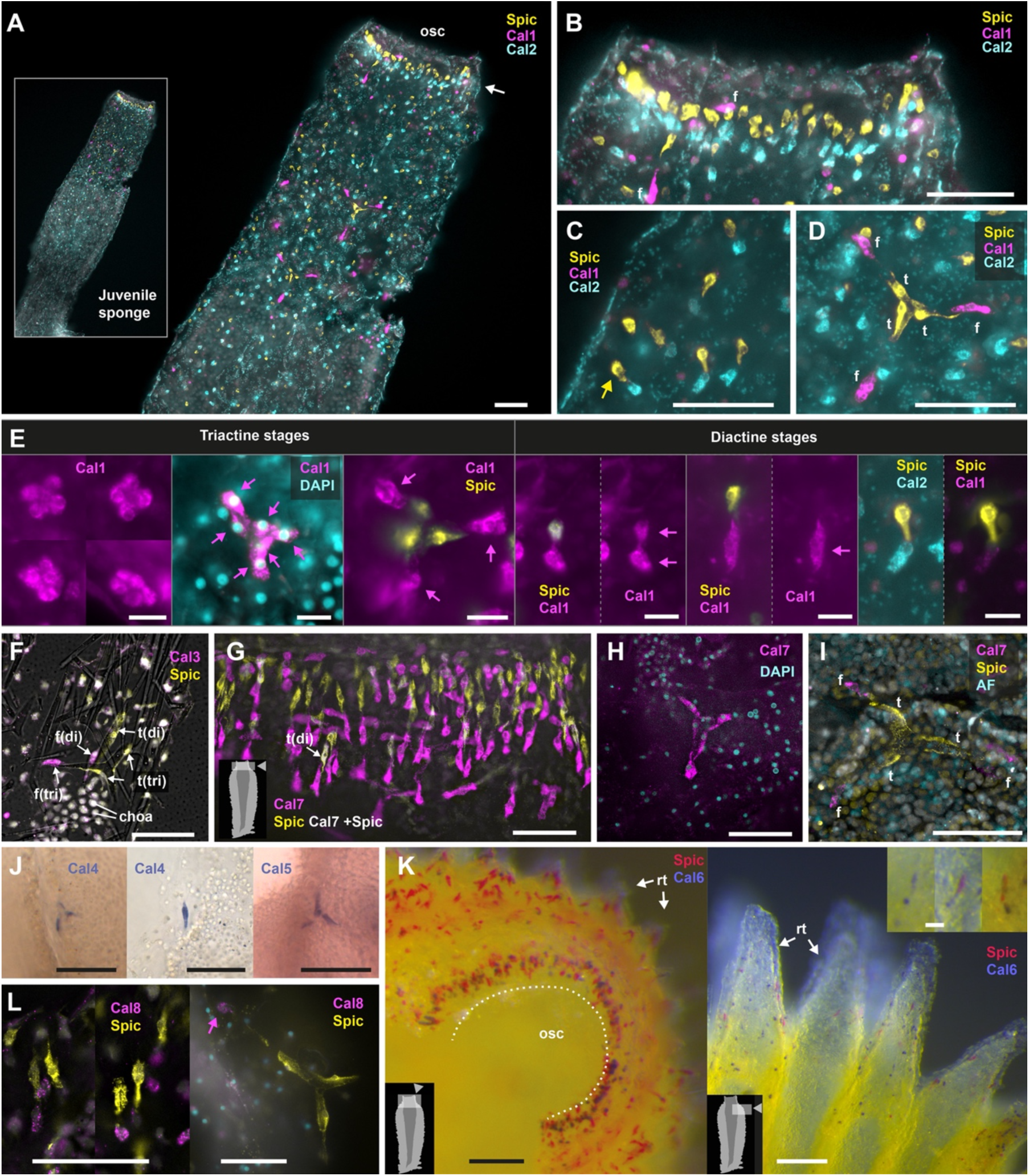
Expression of calcarins. Insets indicate the location of the depicted view within the sponge body, where applicable. A-D: Cal1, Cal2, and Spiculin Expression in a regenerated *Sycon ciliatum* at the asconoid juvenile stage. Scale bars = 50 µm. A: Overview of the entire specimen, highlighting distinct gene expression with minimal co-expression in the apical half of the sponge. B: Detailed view on the expression around the ocular region; Spiculin in sclerocytes’ apical thickener cells, Cal2 in basal founder cells, and Cal1 in triactine/tetractine founder cells. Arrow points to the ring of founder and thickener cells that form the oscular diactines. C: Sponge wall detail; Cal2 in diactine founder cells, Spiculin in thickener cells. D: Triactine/tetractine founder cells expressing Cal1, thickener cells expressing Spiculin. E: Cal1 expression in founder cells ceases when they transform into thickener cells, and Spiculin expression sets in. Cal1 continues to be expressed in actine-producing founder cells in triactines, but in the diactine actine-forming founder cell, it is replaced by Cal 2 expression in later stages. Scale bars 10 µm. F: Expression of Cal3 in founder cells and Spiculin in thickener cells attached to the preserved diactine (di) and triactine (tri) spicules (overlay of light microcopic picture). Note how thickener cells thinly ensheath the spicule. Scale bar: 50 µm. G: Cal7 expression in the founder cells of oscular diactines. Co-expression of Spiculin and Cal7 rarely occurs in transient stages of emerging thickener cells. Scale bar: 50 µm. H: Early triactine stage with six founder cells expressing Cal7. Scale bar: 50 µm. I: In later triactine formation stages, thickener cells no longer express Cal7. Scale bar: 50 µm. J: Expression of Cal4 and Cal5 in thickener cells. Scale bar: 50 µm. K: Cal6 and Spiculin expression in oscular region. Osc: oscular opening, Expression at the distal end of radial tubes of the body wall (rt). Scale bars= 100 µm, inset 20 µm. .L: Expression of Cal8 in founder cells and Spiculin in thickener cells of diactines at the end of radial tubes (left) and a triactine (right). To improve accessibility for individuals with red/green color vision deficiency, original RGB channel colors were modified to a cyan/magenta/blue color scheme. AF: Autofluorescence, osc= osculum, rt: radial tubes, Spic: Spiculin.

Cal1 is predominantly observed in the founder cells of triactines and tetractines and only minimal overlap with Cal2 expression (Fig. 3 A-D). Initially expressed in all six founder cells, expression ceases in the central sclerocytes after they transform into thickener cells. The Cal1 signal persists in the remaining founder cells at the tips of the growing spicules (Fig. 3 D, E). In the founder cells of the diactines, Cal1 is only expressed transiently, as is evident by the lack of Cal1-signal in most oscular diactine founder cells (Fig. 3. A, B). Occasionally, we observed co-expression of Spiculin and Cal1 in diactine sclerocytes, presumably during the onset of the conversion of a diactine founder cell to a thickener cell (Fig. 3, E). At slightly later stages (recognizable by the elongated cell shape), Cal1 is no longer expressed in the thickener cells. At even later stages, Cal1 expression also ceases in the founder cells of the diactines, which instead express Cal2 (Figs. 3 E, S2). The expression of Cal2 and Spiculin can be recognized in two superimposed rings of cells around the osculum, representing the founder cells and the associated thickener cells of the oscular diactines (Fig. 3 B).

Cal3, Cal7, and Cal8 are produced by all founder cells, regardless of the type of spicule they form (Fig. 3 F-I, L). During the transformation into thickener cells, the expression of these Calcarins ceases and is replaced by the expression of Spiculin. Again, co-expression of some of these calcarins with Spiculin was observed in a few cells, probably in sclerocytes transitioning from founder to thickener cells (Fig. 3 G). Cal4 is expressed in thickener cells, Cal5 only in thickener cells of triactines, like Spiculin and Triactinin, respectively (Fig. 3 J). Cal6 expression mirrors that of Cal2, occurring in rounded cells at the distal tip of radial tubes and in a ring of cells around the oscular ring (Fig. 3K). In these regions, Spiculin-expressing cells are spatially associated. In regions without diactines, like the atrial cavity, no Cal6 expression occurs, even if Spiculin expression indicates ongoing production of triactines and tetractines (Fig. S3). Like Cal2, Cal6 must be expressed by later-stage diactine founder cells, which appear more spherical than elongated. At the end of radial tubes, Cal2 and Cal6 positive founder cells are in contact with choanocytes (Fig. S3 A). In summary, calcarin expression in sclerocytes varies across spicule formation stages and between sclerocytes of different spicule types (Fig. 4), adding further complexity to the spatio-temporal expression of biomineralization genes that was reported before (Voigt et al., 2017). Notably, in young coral polyps, normalized and scaled expression data of the raw cell counts of 980 calicoblastic cells indicate non-overlapping expression of two galaxin-like proteins, an uncharacterized skeletal matrix protein and a skeletal-aspartic acid-rich protein (Fig. S4), suggesting spatio-temporal expression changes of skeletal matrix proteins also occurs in calicoblastic cells of corals.

**Figure 4:**
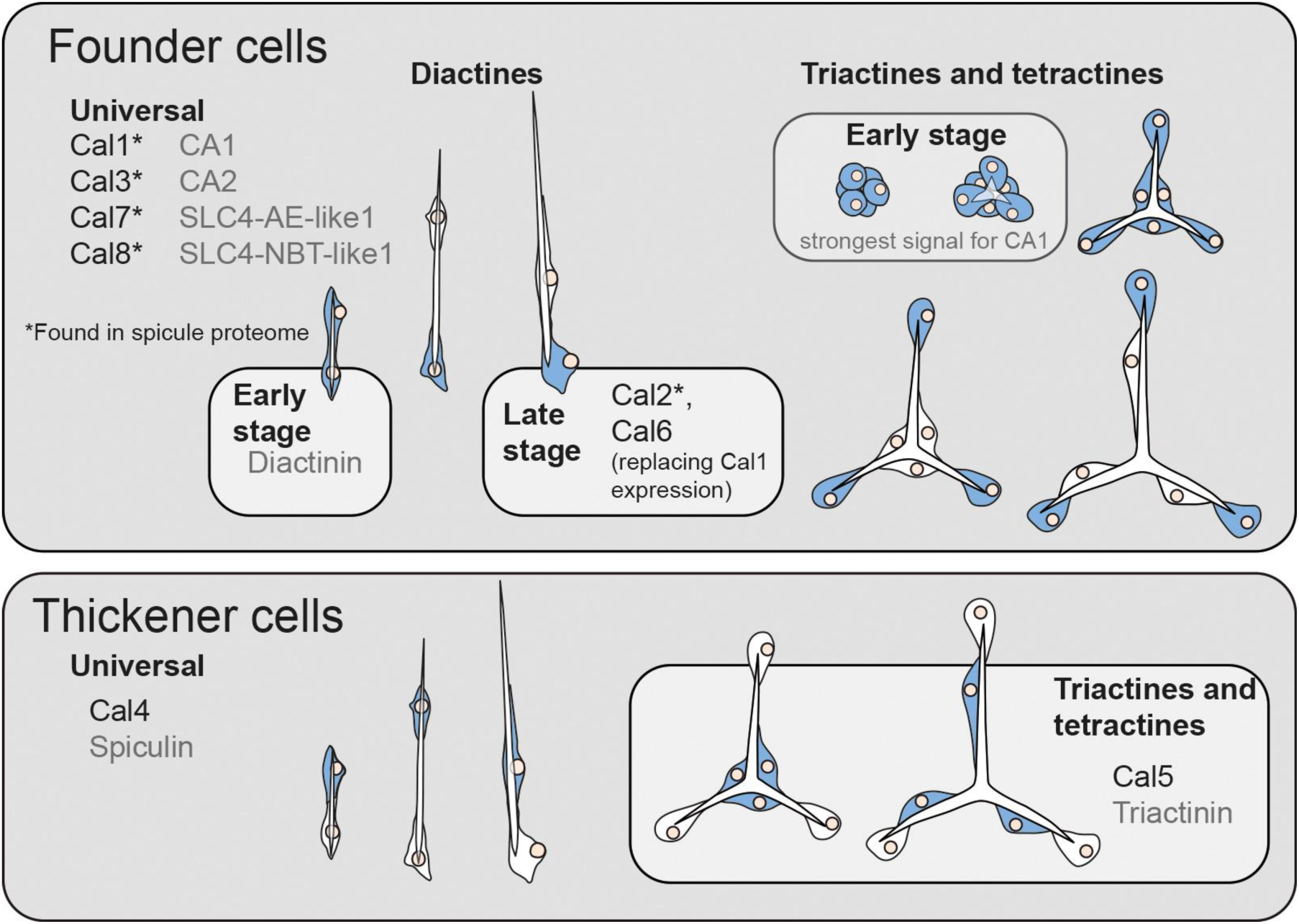
Summary of expression changes of biomineralization genes in sclerocytes (expressing cells in blue). In inital spicule formation stages, all sclerocytes act as founder cells. Genes with expression patterns that were described peviously (Voigt et al., 2017, 2014) are shown in grey.

### Calcarins and other proteins are embedded in the spicule matrix

We extracted spicules from the tissue using sodium hypochlorite to examine the proteins incorporated in the biomineral. The spicules were then washed with water and dissolved in acetic acid. Mass spectrometry was employed to analyze the acid-insoluble proteins, while the acid-soluble ones were too diluted for adequate examination. With the obtained spectra, we identified thirty-five proteins (1.0% FDR protein threshold, with at least two peptides per protein, Dataset S2). Fifteen of these proteins are encoded by over-expressed transcripts in the oscular region (Fig. 5 A). We consider these proteins to be specialized biomineralization proteins. Several calcarins expressed in founder cells (Cal1, Cal2, Cal3, Cal7, Cal8) were most prominent. Additional calcarins of the skeletal matrix were Cal9, Cal11, and Cal10, although the latter was only supported by one peptide.

**Figure 5:**
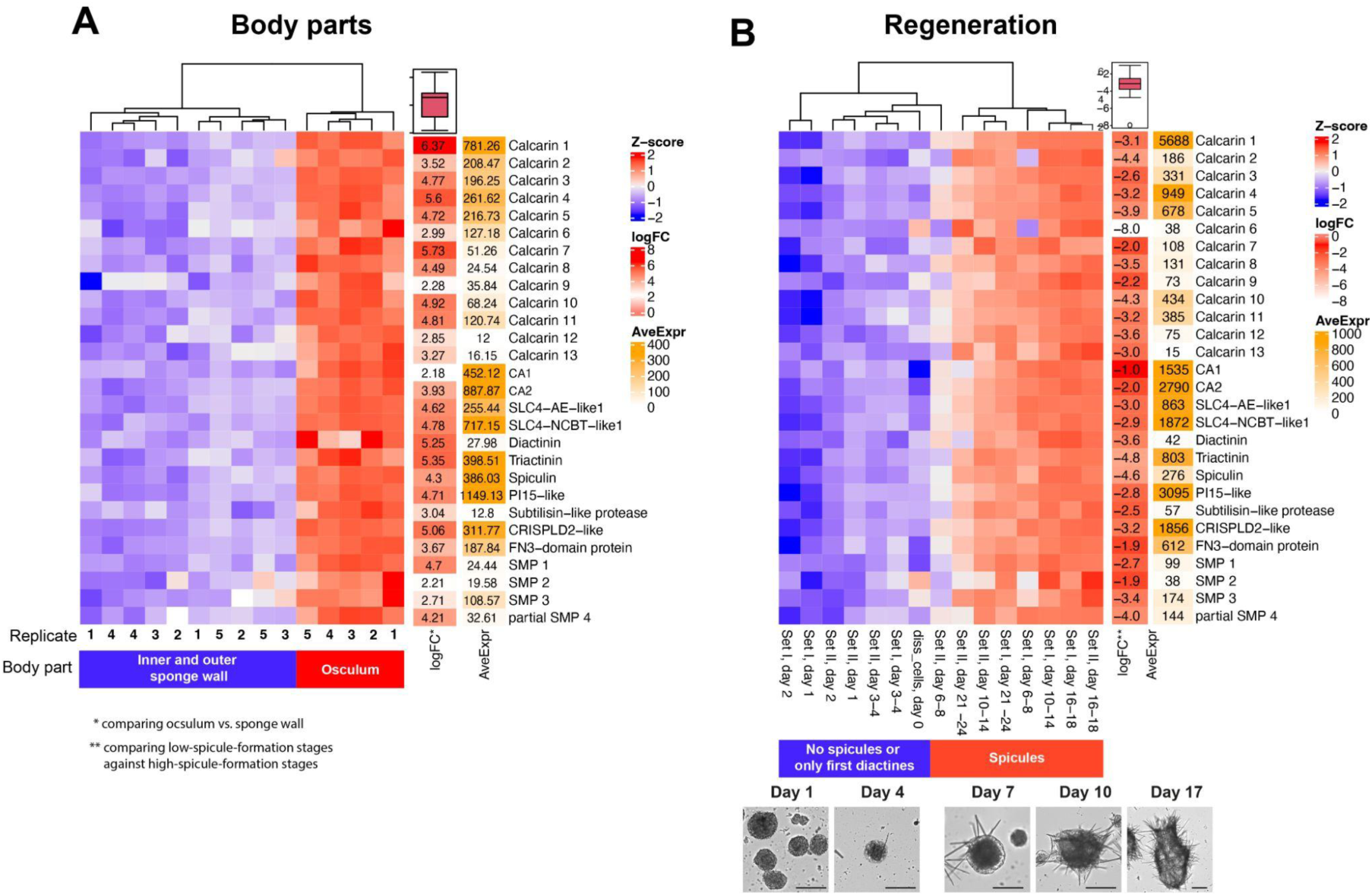
Differential gene expression of 13 calcarins and other confirmed or candidate biomineralization genes. A: Osculum region vs. sponge wall; B: Changes in relative expression during whole-body regeneration.

In addition, we found three secreted proteins without any domain or other prominent features other than a signal peptide, which we refer to as skeletal matrix proteins 1-3 (SMP 1-3). Other spicule matrix proteins contain recognizable domains. Two proteins, similar to Cysteine-rich secretory protein LCCL domain-containing 2-like (CAH0893205) and Peptidase inhibitor 15-like protein (CAH0891878), both belong to the CAP superfamily. Further, we detected a Fibronectin III-domain-containing protein (CAH0869470), a Subtilisin-like protease (CAH0797261), and an EGF-like-domain-containing protein (CAH0821874). Both the LCCL domain-containing 2-like protein and the Fibronectin III-domain-containing protein are acidic with isoelectric points of 4.34 and 4.75, respectively. Thickener-cell-specific proteins, such as Cal4, Cal5, or the Asx-rich proteins Triactinin and Spiculin, were not observed in the spicule matrix.

When comparing the fifteen spicule matrix proteins of *Sycon* to the skeletal matrix proteins of a *Vaceletia* sp. (Germer et al., 2015), a demosponge with a calcium carbonate (aragonite) basal skeleton, their similarity is limited to Subtilisin-like proteases found in both proteomes. Additional similarity can be seen only in eight additional proteins of presumably intracellular origin (annotated as or containing domains of Histones, Actin, Tubulin, Elongation factor 1a, mitochondrial ATP synthase alpha, dataset S3), which were not overexpressed in the oscular region of *Sycon ciliatum*. These proteins may represent ubiquitous extracellular matrix components or, particularly in the case of proteins with intracellular origins, could result from contamination by residual cellular material or their incorporation into the forming biomineral, as suggested before (Ramos-Silva et al. 2013; Germer et al. 2015) and we do not consider them to have specific function in biomineralization.

### Expression of biomineralization genes during whole-body regeneration

Dissociated cells of *Sycon ciliatum* form spherical cell aggregations that differentiate into small functional sponges within 16 to 18 days. This regeneration resembles the post-larval development in *Sycon ciliatum* after the first days (Soubigou et al., 2020). The first stages are free of spicules, and the first diactines appear 3-4 days after reaggregation. Spicule production, including triactines and tetractines, increases in the following days. After about 16 days, an osculum is formed and framed by diactines. Reanalyzing raw RNAseq data covering the complete regeneration process (Table S2) (Soubigou et al., 2020), we find down-regulated expression of calcarins 1-13, the other spicule-matrix proteins, and previously identified sclerocyte-specific genes (Voigt et al., 2017, 2014) in regeneration stages with no or on-setting spicule formation (days 1-4) compared to later stages (days 6-18) with higher numbers of newly forming spicules (Fig. 5 B). Noteworthy, dissociated cells (day 0) also have low expression of these sclerocyte-specific genes, suggesting that most sclerocytes do not survive the cell-dissociation process.

### The majority of calcareous sponge biomineralization genes show concerted changes in expression in different biological settings

We performed a Weighted Gene Co-expression Network Analysis (WGCNA) of the body part and regeneration datasets to identify co-expression modules representing groups of genes displaying concerted expression patterns. The analysis provided eight meta-modules, of which four showed significant changes in expression module eigengenes —summary profiles that capture the overall expression pattern of each module— between samples with high spicule formation context (osculum region and regeneration stages older than four days) and samples with low spicule formation (sponge-wall and early regeneration stages until day 3-4) (Fig. S5). One meta-module “midnightblue” showed higher module eigengene expressions in the context of higher spicule formation. The module includes 196 genes, all differentially expressed in the body-part dataset. Of these, 189 genes were overexpressed in the oscular region, including the suggested biomineralization genes, except for Cal6, Cal9, Cal13, and SMP2. Only seven genes in this meta-module are underexpressed in the oscular region. Some enriched biological process GO-terms of genes in this meta-module could be relevant in the context of spicule formation, e.g., monoatomic anion transport, cell junction organization and assembly, and cell-to-cell signaling (Fig. S6). Eight proteins in the meta module”midnightblue” are involved in the regulation of transcription (GO:0006355), including SciMsx and other transcription factors. The meta module also contains five genes annotated as similar to Integrin alpha 8 or Paxillin from the transforming growth factor beta signaling pathway (GO:0007179), and three genes annotated to belong to the Wnt signaling pathway (GO:0016055), namely SciFzdD, SciDvlB and SciWntI (Dataset S4).

### Calcarins and related galaxin-like proteins in other species

To identify potential homologs of calcarins in a range of other species, we employed Orthofinder to analyze transcriptomes from calcareous sponges, other sponge classes, octocorals, and stony corals (Table S3). This analysis segregated fourteen *Sycon ciliatum* calcarins into distinct orthogroups while grouping Cal2, Cal4, and Cal6 in a single orthogroup. Calcarins in each orthogroup were found in the transcriptomes of other calcareous sponges, either exclusively in Calcaronea (Cal3, Cal5, Cal13, Cal15, and Cal17) or in both Calcaronea and Calcinea (Fig. 2 C, Dataset S5). Within the same orthogroup, Cal2, Cal4, and Cal6 exhibited sequence identities of 25% to 36% among themselves, while each showed a higher match in *Grantia compressa*, with identities ranging from 44% to 58%.

Two orthogroups, one containing Cal2, Cal4, Cal6, and the other Cal15, also included galaxin-like sequences from octocorals. Orthogroups including Cal12 and Cal14 include sequences from the genome of the scleractinian coral *Stylophora pistillata* but not from *Acropora millepora*. Despite their presence in the same orthogroups, the octocoral and stony coral proteins were only distantly related to the calcareous sponge calcarins (e.g., 12-24% identity between octocoral and calcareous sequences in orthogroup Cal 2-4-6), resulting in poor sequence alignment. Alphafold-predictions suggest that the galaxin-like proteins from the octocorals *Heliopora coerulea* and *Tubipora muscia* share a similar structure with Cal2, 4 and 6, each featuring 12 beta-hairpins (Fig. S7). In contrast, galaxin-like proteins from *Stylophora* exhibit a substantially higher number of beta-hairpins than the *Sycon ciliatum* calcarins that are part of the same orthogroup (Fig. S7). Furthermore, none of these scleractinian and octocoral proteins included in the calcarins orthogroup have been directly linked to biomineralization: they were not detected in these species’ skeletons (Conci et al., 2018, Peled et al., 2020) nor have their expression patterns been characterized. Their homology to calcarins, therefore, remains to be determined. Finally, no other sponge proteins grouped with the 17 identified calcarins from *Sycon ciliatum*, although a few galaxin-like sequences were detectable using BLASTp (Dataset S6) —precisely one in Homoscleromorpha, one in each Hexactinellida species, and four in the transcriptome but not the skeletal proteome (Germer et al., 2015) of the demosponge *Vaceletia* sp.. While the galaxin-like transcripts of *Vaceletia* are short and incomplete, the hexactinellid and homoscleromorph proteins are complete and are large proteins of about 5,000 amino acids. A signal peptide suggests that they, too, are secreted. In contrast to calcarins or galaxins, they contain multiple recognizable domains, such as several Fibronectin type III and Laminin EGF domains. Only about 200 amino acids of these proteins are made from the di-cysteine-containing region of the Galaxin BLASTp-hit, which lies between Fibronectin type III domains (Fig. S8). The demosponges *Ephydatia muelleri* and *Tethya wilhelma* lack galaxin-like sequences in their genomes.

Galaxins and galaxin-like proteins are attributed to a PANTHER “family” PTHR34490, and in InterPro (https://www.ebi.ac.uk/interpro/), 623 proteins are currently annotated to this family (with 381 available Alpha-Fold structures, see Dataset S6). They occur in several animal phyla, but also other eukaryotes and archaeans. Species without skeletons, such as the cnidarians *Hydra*, *Actinia*, *Exaiptasia*, and *Nematostella*, also possess galaxin-like proteins assigned to the PANTHER “family” PTHR34490 (Dataset S6).

### Sequential order and expression of biomineralization genes

In the *Sycon ciliatum* genome, the 17 calcarins occur on chromosomes 1, 2, 4, 7, and 13. On chromosome 2, Cal12, Cal4, Cal6, and Cal2 are positioned sequentially (Fig. 6, dataset S7). Cal4, Cal6, and Cal2 belong to a single orthogroup, indicating their homology (Fig. 2). Cal4 is expressed in thickener cells, while Cal2 and Cal6 have identical expression patterns, suggesting they are produced by diactine founder cells during the late diactine formation stage (see above). Cal12 has low expression levels, and we did not locate its expression. The founder-cell specific carbonic anhydrase gene CA2 is situated on the reverse strand of chromosome 4, flanked by CA8 and iCA3 on the forward strand. Upstream on the same chromosome, other membrane-bound carbonic anhydrases (Voigt et al., 2021) are present, including CA4, CA5, CA6, and CA7.

**Figure 6:**
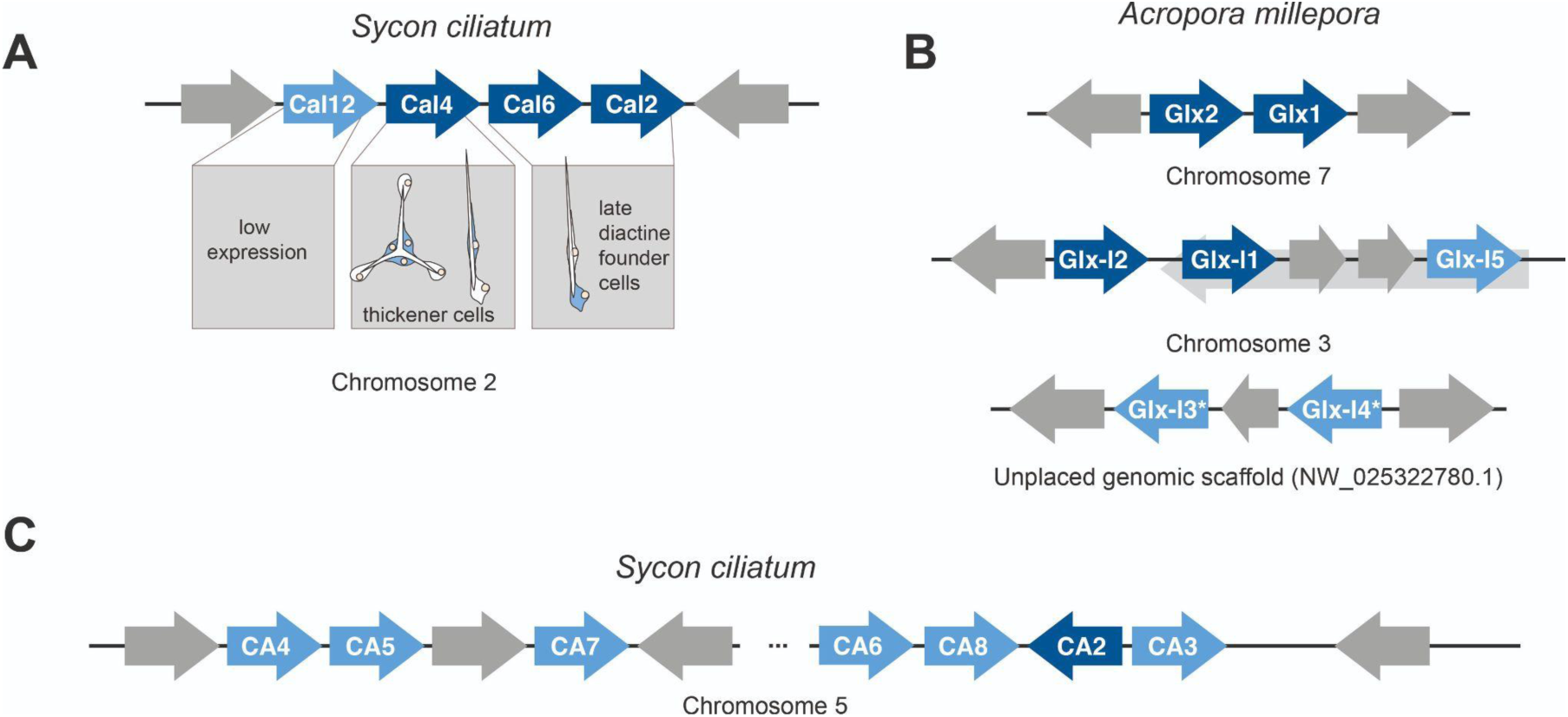
Arrangements of biomineralization genes (dark blue) and related genes (lighter blue). A: Calcarins (Cal) in *Sycon ciliatum*; B: Galaxin (Glx) and Galaxin-like (Glx-l) proteins in the stony coral *Acropora millepora*; C: Membrane-bound carbonic anhydrases (CA) in *Sycon ciliatum*. *predicted nested genes not shown.

In the coral *Acropora millepora*, Galaxin1 and Galaxin2 are located subsequently on the genome, making a past gene duplication likely as well. The galaxin-like proteins 1 and 3 also occur in subsequent pairs or triplets. The genes for Galaxin-like 2, 4, and 5, as well as Galaxin-like 3 and 4, are separated by a maximum of two short predicted genes (Fig. 6).

## Discussion

### Calcarins are galaxin-like biomineralization proteins

Our study significantly expanded the known non-bilaterian biomineralization genes and found that galaxin-like calcarins are key components of the calcareous sponge biomineralization machinery. Most calcarins could be linked to biomineralization by their higher expression levels in body parts or regeneration stages with increased spicule formation, sclerocyte-specific expression, presence in the extracted spicule matrix, or a combination of these. Additionally, they change their expression in concert with other previously characterized biomineralization genes.

In stony corals and octocorals, certain galaxins and galaxin-like proteins are part of the coral skeletal organic matrix (Fukuda et al., 2003; Watanabe et al., 2003) or expressed briefly before the onset of biomineralization in the primary polyp (Reyes-Bermudez et al., 2009). Typically, one or two galaxins occur in stony coral (Peled et al., 2020; Ramos-Silva et al., 2013; Takeuchi et al., 2016) or octocoral (Conci et al., 2020; Le Roy et al., 2021) skeletons. In the spicule matrix of *Sycon ciliatum*, however, we found 7-8 calcarins, so these galaxin-like matrix proteins are much more diverse in calcareous sponges than in corals.

### Role of calcarins, galaxins, and galaxin-like proteins

The temporal and spatial differences in expression patterns of calcarins suggest specialized functions in controlling the biomineralization process. The observed expression patterns of calcarins show that initially, all sclerocytes involved in the formation of a single spicule are functional founder cells expressing the same repertoire of biomineralization genes (e.g., expression of Cal1 and Cal7, Fig. 3 E & H). Founder cell-specific calcarins are integrated into the spicule matrix, much like galaxins and galaxin-like proteins are in coral skeletons. In corals, galaxins and galaxin-like proteins may provide either a structural framework for the growing biomineral or might influence the nucleation and growth of the carbonate crystals (Fukuda et al., 2003; Watanabe et al., 2003), although their function is not fully understood. We assume similar roles for the calcarins we identified from the spicule matrix. A recent study has shown that calcareous sponge spicules grow by addition of calcium carbonate granules that form near the membranes of active sclerocytes (Wendt et al. 2025). Secreted calcarins may play a role in the formation of these granules and are likely among the previously unidentified intercalated proteins that were supposed to influence crystal growth in calcareous sponges by selectively inhibiting growth in specific directions (Aizenberg et al., 1996). We hypothesize that differences in calcarin composition in different spicule types influence their specific crystallographic features and possibly their macromorphology (Aizenberg et al., 1995, 1994).

Our results suggest thickener cells emerge from founder cells by changing gene expression, including calcarins. Thickener-cell-specific calcarins and acidic proteins are not found in the spicule matrix, implying that *Sycon ciliatum* thickener cells do not add significant calcite material to the spicule. Instead, biomineralization genes secreted by thickener cells (Cal4, Cal5, acidic proteins Triactinin and Spiculin) may affect only the spicule’s surface. For instance, spicules in one Calcinea species have a calcitic core surrounded by amorphous calcium carbonate cored again by a thin calcitic sheath (Aizenberg et al., 2003). Here, acidic proteins and calcarins might be involved in transforming amorphous calcium carbonate to calcite, but due to the thin nature of the sheath, they are more likely to be lost (due to dissolution during clean-up) or remain below the detection threshold.

### The calcareous sponge biomineralization toolkit resembles that of corals

Our widened understanding of the calcareous sponge biomineralization machinery revealed that the identified effector genes are distinct from those reported from other sponges. This is not surprising because even the formation of siliceous spicules in the classes Demospongiae and Hexactinellida show little overlap (Francis et al., 2023; Shimizu et al., 2024), which indicates that the formation of spicules in these three extant sponge groups evolved independently. Based on our data, we can also exclude that there are major genetic similarities between the formation of calcite spicules of calcareous sponges and the aragonitic basal skeletons of some demosponges, called sclerosponges. Only specific subtilisin-like protease and carbonic anhydrases are shared in the biomineralization machinery of the sclerosponge *Vaceletia* and the calcarean *Sycon ciliatum*. Both belong to larger gene families, and phylogenetic analyses of carbonic anhydrases suggests that they were independently recruited for biomineralization in calcareous sponges and carbonate producing demosponges (Voigt et al., 2021, 2014). Spherulin, a skeletal protein of aragonitic sclerosponges (Germer et al., 2015; Jackson et al., 2011), is not present in the calcareous sponge genome. Although we identified Galaxin-like sequences in *Vaceletia’s* transcriptome, the proteins are not reported from its skeleton (Germer et al., 2015). In contrast, the galaxin-like calcarins are diverse in calcareous sponges and their spicules.

Considering these genetic differences in sponge biomineralization between taxa that secrete aragonite (*Astrosclera*, *Vaceletia*) and calcite (*Sycon*), it is unexpected that the essential biomineralization effector genes of calcareous sponges that produce calcite resemble those utilized by aragonitic stony corals, suggesting parallel evolution in gene recruitment between these calcifying organisms. The gene families involved are key parts of the pre-adapted carbonate “biomineralization toolkit” of their common ancestor, including essential pH regulators and for the directional transport of inorganic carbon, i.e., specialized carbonic anhydrases (Bertucci et al., 2013; Voigt et al., 2014) and bicarbonate transporters (Voigt et al., 2017; Zoccola et al., 2015) (Fig. S9. Galaxin and galaxin-like proteins from corals and calcarins from calcareous sponges extend the set of similar biomineralization genes.

### Possible origin of galaxins, galaxin-like proteins and calcarins

The high sequence divergence within calcarins, galaxin, and galaxin-like proteins makes a resolved phylogenetic analysis of all these genes impossible. The likeliest origin of these biomineralization proteins is the recruitment of secreted di-cysteine-rich proteins with other functions and probably different origins. The characteristic feature of these proteins is the more or less evenly distributed dicysteines, which, according to the Alpha-Fold prediction, establish a similar tertiary structure with beta-hairpins connected via disulfide bridges. Secreted proteins with this structure are not limited to calcifying organisms and can be portions of larger proteins. For example, in hexactinellid and homoscleromorph sponges, the few proteins similar to calcarins are large and contain numerous functional domains, while the typical di-cysteine region only resembles a small proportion of the protein. Clearly, then, galaxin-like proteins in non-calcifying organisms have other functions.

Although our results clearly show that some calcarins have a common origin, it is unclear whether galaxin-like proteins generally have common or multiple convergent origins. For example, the di-cysteine regions of galaxin and galaxin-like proteins of the Scleractinia consist of more or less conserved repeat motifs of different lengths (Fukuda et al., 2003; Reyes-Bermudez et al., 2009). These galaxin-like proteins may result from repeated duplication of an original motif, gradually extending the di-cysteine region. In contrast to the galaxin-like proteins of corals, the calcareous sponge calcarins do not show such detailed repeat structures, which could indicate a different origin.

### Gene duplication gave rise to and diversified biomineralization genes in calcareous sponges and corals

Our results provide evidence about the origin and diversification of biomineralization genes. The similarity and sequential arrangement of Cal4, Cal6, and Cal2 on chromosome 2 in the *Sycon ciliatum* genome suggest that these genes originated from two successive gene duplications.

Their different expression patterns reveal that a function change must have occurred after the gene duplication. Accordingly, the first duplication event produced two copies, leading to one gene specialized for thickener cells and another for late diactine founder cells. Subsequently, the latter gene duplicated again, resulting in the origin of Cal2 and Cal6, which, despite beginning to diverge in sequence, maintain identical expression patterns. This sequence of events highlights the role of gene duplication in the functional diversification of calcarin genes in *Sycon ciliatum*.

Similar observations for galaxins and galaxin-like genes of the stony coral *Acropora millepora* suggest that gene duplication is a typical pattern in galaxins and galaxin-like proteins. Galaxin1 and Galaxin2 also occur adjacently in the coral’s genome, indicating past gene duplication. The clustering of galaxin-like proteins in pairs or triplets and the separation of genes for Galaxin-like2, 4, and 5, as well as Galaxin-like3 and 4, by a maximum of two short predicted genes, may also be evident for their origin by past duplication events. The fact that one of the *Sycon ciliatum* sclerocyte-specific carbonic anhydrases, SciCA2, is likewise located on the reverse strand of chromosome 4 and flanked by two non-biomineralizing carbonic anhydrases (SciCA8 and SciCA3) (Voigt et al., 2014) encoded on the forward strand provides additional support that gene duplication followed by a neofunctionalization of SciCA2 is the origin of this biomineralization gene. A similar process, tandem gene duplication, neofunctionalization, and inversion, gave rise to the coral-specific biomineralization-specific bicarbonate transporter SLC4γ (Tinoco et al., 2023). Our data show that similar genes in both lineages were independently duplicated, and one copy was recruited in the biomineralization machinery by a functional shift and expression in calcifying cells. The example of the three calcarins provides evidence of how the biomineralization process can evolve to become more and more finetuned by further gene duplications and subfunctionalization, e.g., in our case, becoming specific for a distinct spicule type.

### Gene regulatory networks

While our study identified biomineralization genes at the effector level, more research is required to understand the underlying gene regulatory networks, e.g., what triggers the expression changes in some of the spicule’s sclerocytes to change from functioning as a founder cell to a thickener cell. The genes that change expression in meta-module midnightblue might include some key players, as they are enriched in some regulatory biological process GO-terms (Fig. S6). Cell-cell signaling between the cooperating sclerocytes could play a critical role in controlling the temporal change in gene expression, and several apically overexpressed genes reported here may be involved, such as components of the Wnt or BMP signaling pathways, involved in mammal bone morphogenesis and homeostasis (Sánchez-Duffhues et al., 2015; Zhong et al., 2014). However, what these are and how the control mechanisms function must be investigated in further studies because, in both biological scenarios, apical gene expression or regeneration, additional growth, and differentiation processes run parallel to spicule formation, which we cannot differentiate from each other with the methods used here.

## Conclusion

We have identified calcarins, galaxin-like proteins, as novel components of the biomineralization toolkit in calcareous sponges. The diversity of calcarins in the calcitic sponge spicules, compared to galaxin and galaxin-like proteins in aragonitic coral skeletons, underscores their specialized roles in biomineralization. Their varied spatial and temporal expression patterns highlight the precise genetic regulation involved in calcification in this class of sponges. Our results show that similar genes or genes from the same families surprisingly are involved in biomineralization in both calcareous sponges and corals that produce different calcium carbonate polymorphs, calcite vs. aragonite, respectively. This demonstrates that there are parallels between the formation of calcareous sponge spicules and coral skeleton formation at the proteome level. Furthermore, our results highlight how gene duplication and neofunctionalization of an original gene pool gave rise to dedicated biomineralization genes that could further differentiate to refine the biological control of the process. Single-cell expression data of calicoblastic cells in corals indicates that coral biomineralization may equally depend on small-scale spatial and temporal expression changes to finetune calcification, an aspect that requires further study and could help predict the responses of these reef-builders to global environmental changes.

## Materials and Methods

### Specimens, regeneration, and RNASeq of body parts

Specimens of *Sycon ciliatum* were obtained from the AWI Biologische Anstalt Helgoland and sent alive to Munich. Here, specimens were transferred to glass Petri dishes and maintained in seawater for a few days with daily seawater changes. To verify the timing of the occurrence of spicules during the regeneration of *Sycon ciliatum*, we performed a regeneration experiment of the *Sycon ciliatum* as used for the regeneration RNA-Seq dataset (Soubigou et al., 2020). We mechanically dissociated sponge cells through a 60 µm strainer, then incubated the cells in filtered seawater at 14°C in Petri dishes, changing the water every 1-2 days. The timing of whole-body regeneration stages and occurrence of spicules aligned with prior studies. After 30 days, the experiment ended, and we fixed some juvenile asconoid sponges for *in situ* hybridization.

For DGE analysis, we dissected three body parts from five living specimens (Fig. 1): (1) The oscular region with mainly large diactine and tetractine spicules, (2) inner sponge wall, including the atrial skeleton (tetractines + diactines) and the proximal radial tube (triactines), (3) outer sponge wall, including the distal parts of the radial tube with triactines and tufts of curved diactines.

The body parts were instantly transferred into the lysis buffer of the RNA-Duet extraction kit (Zymo). We extracted RNA according to the manufacturer’s instructions and confirmed its integrity with an Agilent Bioanalyzer using the RNA 6000 Nano Kit. We used the SENSE mRNA-Seq Library Prep Kit V2 for Illumina (Lexogen) to produce sequencing libraries. The fifteen libraries were pooled and sequenced on a HiSeq Illumina Sequencer at the Gene Center of LMU. 100 bp paired raw reads were quality-checked with FastQC and trimmed to 92 bp to remove low-quality 3’ bases.

### Differential gene expression analysis

Trimmed raw reads (Table S1) were mapped to a high-quality transcriptome of *Sycon ciliatum* (Caglar et al., 2021) using Geneious (Kearse, M., Moir, R., Wilson, A., Stones-Havas, S., Cheung, M., Sturrock, S., Buxton, S., Cooper, A., Markowitz, S., Duran, C., Thierer, T., Ashton, B., Mentjies, P., & Drummond, A., 2012). We omitted non-mapping reads to remove commensal sequences common in sponge tissues/samples. The trimmed and filtered reads were submitted to the European Nucleotide archive (PRJEB78728). Raw reads from a whole-body regeneration experiment (Soubigou et al., 2020) were downloaded from the European Nucleotide Archive and processed identically (see Table S2 for accession numbers). For mapping, we used the *Sycon ciliatum* transcriptome published in (Caglar et al., 2021). This transcriptome showed better BUSCO (Manni et al., 2021) values than a transcriptome assembled using Trinity and the filtered reads from our experiments and was therefore prefered for mapping prior to DGE analysis with DESeq2 (Love et al., 2014). Gene and transcript counts for each filtered set were obtained with salmon (Patro et al., 2017) and combined into count matrices for the body parts experiment and the regeneration data. For the body-part dataset, we compared gene expression of the apical osculum region with increased spicule formation with gene expression of the more basally located regions of the sponge wall (inner and outer side). In the regeneration data set, we identified the differentially expressed genes for two regeneration series between the initial spicule-free stages (day 1+2) and the subsequent stages that produce spicules (days 3-24). Differentially expressed genes (|log2-fold change| ≥ 2, padj < 0.01) were identified, extracted from the reference transcriptome, and used in subsequent analysis.

### WGCNA

We combined both count datasets, filtered them to exclude low-count genes (genes with less than 10 samples with counts >10), and used them to construct a DESeq2 data set in R, followed by variance stabilizing transformation (VST) for normalization. We performed a Weighted Gene Co-expression Network Analysis (WGCNA) (Langfelder and Horvath, 2012, 2008) to identify gene modules associated with spicule formation. For this purpose, a distinction was made between “low spicule formation” for the body wall and the first stages of regeneration with no or only a few first diactines (day 0-4) compared to the other states (high spicule formation). The resulting modules were detected using the dynamic tree cut algorithm, and module eigengenes were calculated. A permutation test (n=1000) was conducted to assess the significance of modules regarding their association with conditions (low vs high spicule formation).

### GO-term enrichment analysis

Best hit proteins with e-values ≤ e-05 were obtained by running BLASTx of *Sycon ciliatum* transcripts (Caglar et al., 2021) against the UniProt database. GO terms for these best hit proteins (e-values ≤ e-05) were retrieved from QuickGO (www.ebi.ac.uk/QuickGO/) using a custom Perl script. Transcript-associated GO terms were compiled by genes and used as input for GO-term enrichment analysis with topGO (Alexa A, 2023) in R. This analysis was performed for genes overexpressed in the osculum region and genes included in the “midnightblue” module identified by WGCNA. GO terms with fewer than 10 annotated genes were excluded. For each ontology (Molecular Function, Biological Process, and Cellular Component), gene lists were mapped to GO terms, and enrichment was tested using Fisher’s exact test with the “classic” algorithm. Significantly enriched GO terms (p ≤ 0.05) were summarized using Revigo (Supek et al., 2011) to provide representative terms. All scripts, input files, and parameters used in the analysis are provided in a GitHub repository.

### OrthoFinder and BLASTp

Raw RNASeq data of several non-bilaterians was downloaded from SRA and reassembled using the TransPi pipeline (Rivera-Vicéns et al., 2022). The predicted proteins of these assemblies, of genomes and transcriptomes of calcareous sponges, sponges from other classes, octocorals, and stony corals were used as input to OrthoFinder to generate orthogroups (Emms and Kelly, 2019), focusing on species in which biomineralization genes are best studied: *Acropra millepora* (Ramos-Silva et al., 2013; Reyes-Bermudez et al., 2009; Takeuchi et al., 2016), *Stylophora pistillata* (Drake et al., 2013; Peled et al., 2020; Zoccola et al., 2015), three octocoral species (Conci et al., 2020), the calcifying demosponge *Vaceletia* sp. (Germer et al., 2015), as well as some other high-quality sponge genomes (Francis et al., 2023, 2017; Kenny et al., 2020; Shimizu et al., 2024). Sequences of calcarins of Calcaronea were extracted from orthogroup sequences and aligned using Muscle (Edgar, 2004). We used BLASTp (Camacho et al., 2009) with *Acropora millepora* Galaxin (D9IQ16) with a maximum E-value of 1e-5 as the threshold to identify proteins similar to galaxins in the proteomes (Dataset S6).

### RNA *in situ* hybridization

We conducted RNA *in situ* hybridization (ISH) to study the spatial and temporal expression of selected genes using standard chromogenic ISH and Hairpin Chain Reaction fluorescent *in situ* hybridization (HCR-FISH). Fixation of specimens and chromogenic ISH followed published procedures (Fortunato et al., 2012). Further details about fixation, probe preparation, and HCR-FISH are provided in the Supplementary Information.

### Identification of spicules organic matrix proteins

Identification and analysis of organic matrix proteins from *Sycon ciliatum* spicules involved isolating 6 g of spicules with a solution of sodium hypochlorite, decalcifying them with acetic acid, and separating the acid-soluble and insoluble fractions, of which only the latter provided enough protein to be further analyzed. Proteomic analysis included gel electrophoresis, in-gel digestion, and LC-MS/MS. Proteins were identified using MASCOT (Perkins et al., 1999) with the predicted proteins from the *Sycon* transcriptome (PRJEB49276), and additional analyses were performed in Scaffold V5.01 (Proteome Software Inc., Portland, USA). Detailed experimental procedures are available in the Supplementary Information.

### Genome analysis

We accessed the assembly of the *Sycon ciliatum* genome (GCA_964019385) of the Tree of Life Programme and (https://www.sanger.ac.uk/programme/tree-of-life/, https://www.sanger.ac.uk/collaboration/aquatic-symbiosis-genomics-project/). Because we found the provided gene predictions were incomplete, we performed gene predictions using BRAKER3 installed on a public Galaxy server installation (https://usegalaxy.eu), using a HiSat2 (Kim et al., 2019) mapping of our filtered reads and the peptides from genomic scaffolds (Fortunato et al., 2014) as training data. For the stony corals, we obtained the annotated genomes from GenBank (*Acropora millepora* v2.1, GCF_013753865, *Stylophora pistillata* v1.1, GCF_002571385). The genetic locations of biomineralization genes, additional carbonic anhydrases, and SLC4 transporters were identified using BlastP. These coordinates were then extracted from the genome’s GFF file.

### Structure predictions of Calcarins

Alphafold express (https://www.line-d.net/alphafold-express) was used to predict the structures of selected calcarins. Alpha-Fold predictions of *Acropora millepora* Galaxin (D9IQ16) and Galaxin 2 (B8UU51) were downloaded from UniProt (UniProt Consortium, 2021) as pdb files. Model confidence for the structure predictions of the beta-hairpin structure was very high (pLDDT >90) in most cases, but it was generally low or lower for adjacent regions. We visualized predicted structures with the Mol* 3D viewer (Sehnal et al., 2021).

### Inspection of skeletal matrix protein expression in coral calicoblasts

To investigate fine-tuned changes in expression occurring in the coral *Stylophora pistillata* calicoblasts, we examined publicly available single-cell sequencing datasets (Levy et al., 2021). Using the species’ online cell atlas tool (https://sebe-lab.shinyapps.io/Stylophora_cell_atlas/), we visualized the expression of skeletal matrix proteins (Peled et al., 2020). This analysis revealed specific expression of 14 skeletal matrix proteins in calicoblast metacells of young coral polyps (Fig. S4), whereas only very few adult coral calicoblast metacells expressed documented skeletal matrix proteins, indicating no or limited calcification activity (Fig. S10). The raw UMI counts (Spis_polyp_sc_UMI_counts.RDS, Spis_adult_sc_UMI_counts.RDS) and the celltype assignments (Spis_polyp_cell_type_assignments.txt, Spis_coral_cell_type_assignments.txt) were downloded from github (https://github.com/sebepedroslab/Stylophora_single_cell_atlas). Single-cell RNA sequencing data from cells expressing more than 100 genes were normalized and scaled using the LogNormalize and ScaleData functions of the Seurat R package version 5.1.0 (Hao et al., 2024), respectively, to adjust gene expression counts to 10,000 molecules per cell, and to center and scale each gene to a mean of zero and a variance of one. Normalized and scaled expression data of calicoblasts were visualized as heatmaps using pheatmap version 1.0.12.

### Data accessibility

Scripts and input files for the analysis are available in a GitHub repository (https://github.com/PalMuc/CalcBiomin/tree/v0.1), and have been archived within a Zenodo repository (https://zenodo.org/records/13847772) *De novo* gene predictions, detailed outputs from Alpha-Fold and the OrthoFinder-Analysis are provided along with a scaffold file for the proteins identified from the spicule matrix and can be accessed via Zenodo (https://zenodo.org/records/14755899).

## Acknowledgments

We gratefully acknowledge the financial support provided by the German Research Foundation (DFG, project VO 2238/1-1). We also thank the Darwin Tree of Life project (https://www.darwintreeoflife.org) and the Aquatic Symbiosis Genome Project (https://www.sanger.ac.uk/collaboration/aquatic-symbiosis-genomics-project/) for sequencing and providing the genomic resources of *Sycon ciliatum*. Some of the calculations were performed on the Galaxy server, which is partially funded by the Collaborative Research Centre 992 Medical Epigenetics (DFG grant SFB 992/1 2012) and the German Federal Ministry of Education and Research (BMBF grants 031 A538A/A538C RBC, 031L0101B/031L0101C de.NBI-epi, and 031L0106 de.STAIR (de.NBI)). We also thank Helmut Blum and Stefan Krebs from the Gene Center (Ludwig-Maximilians-Universität München) for performing the RNASeq sequencing. Special thanks to Nora Dotzler and Lara Aust for their assistance with the HCR-FISH experiments and to Lara Aust for her work on the regeneration of *Sycon ciliatum*. Additionally, we appreciate the help of Bernhard Ruthensteiner (Zoologische Staatssammlung München) with the µCT scans. English language was improved with the help of ChatGTP-4.0 (OpenAI; accessible at https://openai.com/blog/chatgpt).

## Author Contributions

Oliver Voigt: Conception, obtaining funding, data generation, analysis, writing the manuscript Lena Wilde, Fröhlich: Data generation and analysis (proteome)

Benedetta Fradusco: Data generation: RNA extraction, library preparation

Sergio Vargas: Support in data analysis, providing predicted proteins of transcriptomes Gert Wörheide: Providing infrastructure, conception, manuscript editing

All authors have read the final version of the manuscript.

## Supporting Information

### Extended Methods

#### Chromogenic and HCR RNA in situ hybridization

We investigated the spatial and temporal expression of selected calcarins and previously characterized biomineralization genes through standard chromogenic RNA *in situ* hybridization (ISH) and Hairpin Chain Reaction fluorescent *in situ* hybridization (HCR-FISH). Living *Sycon ciliatum* were fixed using MOPS fixation buffer (100 mM MOPS, ph 7.5; 0.5 M sodium chloride; 2 mM MgSO_4_; 4% paraformaldehyde, 0.05% glutaraldehyde) at four °C overnight, followed by dehydration in 70% ethanol and storage at −20°C. Target genes were amplified from cDNA using gene-specific primers (Table S1), and cloned into pCR®4-TOPO® cloning vector (Invitrogen) with T3 or T7 initiation sequence adjacent to the insertion site. The plasmid was sequenced with vector-specific primers to determine the insert direction. From the plasmid, a template for *in vitro* transcription was generated by PCR with the gene-specific forward primer and a vector-specific primer (depending on the insertion direction). Antisense RNA probes for chromogenic ISH were generated by *in vitro* transcription using T3 or T7 RNA polymerase from plasmids or PCR products of the target genes (Table S4). Probes were labeled with digoxigenin or fluorescein with a corresponding RNA labeling kit (Roche). Additional antisense RNA probes were available from a previous study (Voigt et al., 2017). The procedures for ISH followed published protocols (Fortunato et al., 2012; Voigt et al., 2017). We employed a two-probe detection system for double ISH, using NBT/BCIP (Roche) for the first probe and FastRed (Roche) for the second, the documentation was performed using Leica microscopes (M165F, DMLB).

For HCR-FISH, gene-specific probe sets, each consisting of 20 pairs of gene-specific probes (Table S5) based on the target gene’s transcript sequence, were designed and delivered by Molecular Instruments (https://www.molecularinstruments.com), which also delivered corresponding amplifiers, labeled with AF488, AF546, AF594 or AF647. HCR FISH allowed us to visualize up to three target genes with different fluorochromes simultaneously. Tissue preparation for HCR-FISH mirrored the initial steps of ISH up to re-fixation and the subsequent washing steps, followed by whole-mount HCR-FISH as specified by the manufacturer. Post-HCR-FISH, tissues were left in 5x SSC overnight to dissolve remaining spicules that impede probe visualization due to their optical properties. We then mounted the tissue using EverBbrite Hardset (Biotum) for visualization or embedded in Technovit (Kulzer) for sectioning. We used a LEICA Thunder Imager for fluorescent microscopy and photo documentation, using Z-stacking to extend the focal range. Background autofluorescence in thicker tissues was reduced using Leica’s Large Volume Computational Image Clearing (LVCC) implemented in Leica Application Suite (Leica). To improve accessibility for individuals with red/green color vision deficiency, the resulting HCR-FISH images of Fig. 3 were processed in ImageJ by splitting channels, applying cyan, magenta, or yellow lookup tables, converting each channel image to a RGB type image, and combining them using the Z-Project function with maximum intensity projection. A version with the original channel information preserved is provided in Fig. S11.

#### Identification of organic matrix proteins from spicules

*Sycon ciliatum* spicules were isolated by dissolving the surrounding tissue in a 5% sodium hypochlorite solution, followed by several rinses with MiliQ water and air drying. Approximately 6 g of dried spicules were then decalcified overnight in 10% acetic acid at room temperature on an orbital shaker. This process separated the spicules into acid-soluble (ASM) and acid-insoluble matrices (AIM), then divided via centrifugation at 14,000 x g for 30 minutes. The ASM fraction was desalted using Amicon Ultrafiltration devices, and the AIM residues were thoroughly washed, and both fractions were subsequently stored at −20°C.

To prepare samples for LC-MS/MS, they were unthawed, briefly vortexed, and allowed to settle for 3-5 minutes. 250 µl of the semi-transparent phase was then transferred into Eppendorf tubes and centrifuged for 30 s at 5,000 rpm and 4°C. The resulting pellet was washed twice with ultrapure mass-spectrometry grade water (MS-H2O, Merck) and resuspended in 100 µl MS-H_2_O. After sonication with a cup resonator (Bandelin), 2X Laemmli buffer was added, and the samples were incubated at 95°C for 5 minutes, followed by a second sonication. The prepared samples were vortexed and loaded onto a NuPAGE 4-12% Bis-Tris Gel (Invitrogen, USA). Electrophoresis was performed at 200 V until the gel pockets were empty, and the gel was subsequently stained overnight with Roti Blue staining solution (Roth, Germany). Protein-containing gel segments were excised, destained, and washed with 50 mM NH_4_HCO_3_. The supernatant was discarded, and the gel pieces were reduced with 45 mM DTE for 30 minutes at 55°C, then alkylated with 100 mM iodoacetamide in the dark at room temperature. Following alkylation, the gel pieces were washed again, and a sequential in-gel digestion was executed using Lysyl Endopeptidase (Lys-C, 4 hours at 37°C, Mass Spectrometry Grade, FUJIFILM Wako Pure Chemical Corporation, USA) followed by an overnight digestion with 100 ng of sequencing grade modified trypsin (Promega, Germany) at 37°C. The resulting supernatants containing peptides were collected, and the peptides were extracted using 70% acetonitrile (ACN). These collected supernatants were pooled and then dried using a vacuum centrifuge (Bachofer, Germany).

LC-MS/MS analysis was performed using an Ultimate 3000 RSLC (Thermo Fisher Scientific) connected to a Q Exactive HF-X mass spectrometer (Thermo Fisher Scientific). Protein samples were loaded onto trap columns (PepMap 100 C18, 100 µm × 2 cm, 5 µM particles, Thermo Scientific) using a flow rate of 5 µL/min (mobile phase: 0.1% formic acid and 1% acetonitrile in water). Liquid chromatography was performed with an EASY-Spray column (PepMap RSLC C18, 75 µm × 50 cm, 2 µm particles, Thermo Scientific) and a 250 nL/min flow rate. Peptide samples were separated with a two-step gradient from 3% mobile phase B (0.1% formic acid in acetonitrile) to 25% mobile phase B in 30 min, followed by a ramp to 40% for 5 min (mobile phase A: 0.1% formic acid in water). The mass spectrometer was set to data-dependent acquisition mode, performing a maximum of 15 MS/MS spectra per survey scan.

MS/MS spectra were analyzed with MASCOT V2.6.2 (Matrix Science Limited, UK) (Perkins et al., 1999). The predicted proteins from the *Sycon ciliatum* transcriptome (PRJEB49276) served as a database to which we added a predicted protein sequence of Cal8 that we found as a second, untranslated protein encoded on transcript HBWS01158424. We further evaluated the results in Scaffold (v. 5.0.1, Proteome Software Inc., Portland, USA), setting the filters to a protein threshold of 1.0% FDR, the minimum number of peptides =2, and a peptide threshold of 1.0% FDR. The mass spectrometry proteomics data have been deposited to the ProteomeXchange Consortium (http://proteomecentral.proteomexchange.org) via the PRIDE partner repository (Perez-Riverol et al., 2019) with the project accession PXD060105.

#### Legends Datasets S1 to S7

**Dataset S1 (separate file).** Biological Process GO-terms enriched in genes overexpressed in the oscular region of *Sycon ciliatum* (representative terms obtained from Revigo).

**Dataset S2 (separate file).** Proteins identified in the calcareous sponge spicule matrix. Proteins that are overexpressed in the oscular region (2logfold change) are in bold, Calcarins highlighted by yellow background.

**Dataset S3 (separate file)**: *Vaceletia* sp. skeletal proteins similar to *Sycon ciliatum* spicule matrix proteins.

**Dataset S4 (separate file)**. Genes overexpressed in the oscular region and included in meta module midnightblue with selected gene regulatory GO-annotations.

**Dataset S5 (separate file)**. OrthoFinder results for orthogroups that include calcarins and other biomineralization genes.

**Dataset S6 (separate file).** Galaxin BlastP hits in the proteomes used in OrthoFinder analysis and proteins and structures of PANTHER family “PTHR34490”.

**Dataset S7 (separate file).** Genomic position of biomineralization genes in *Sycon ciliatum*, *Acropora millepora*, and *Stylophora pistillata*.

**Fig. S1.**
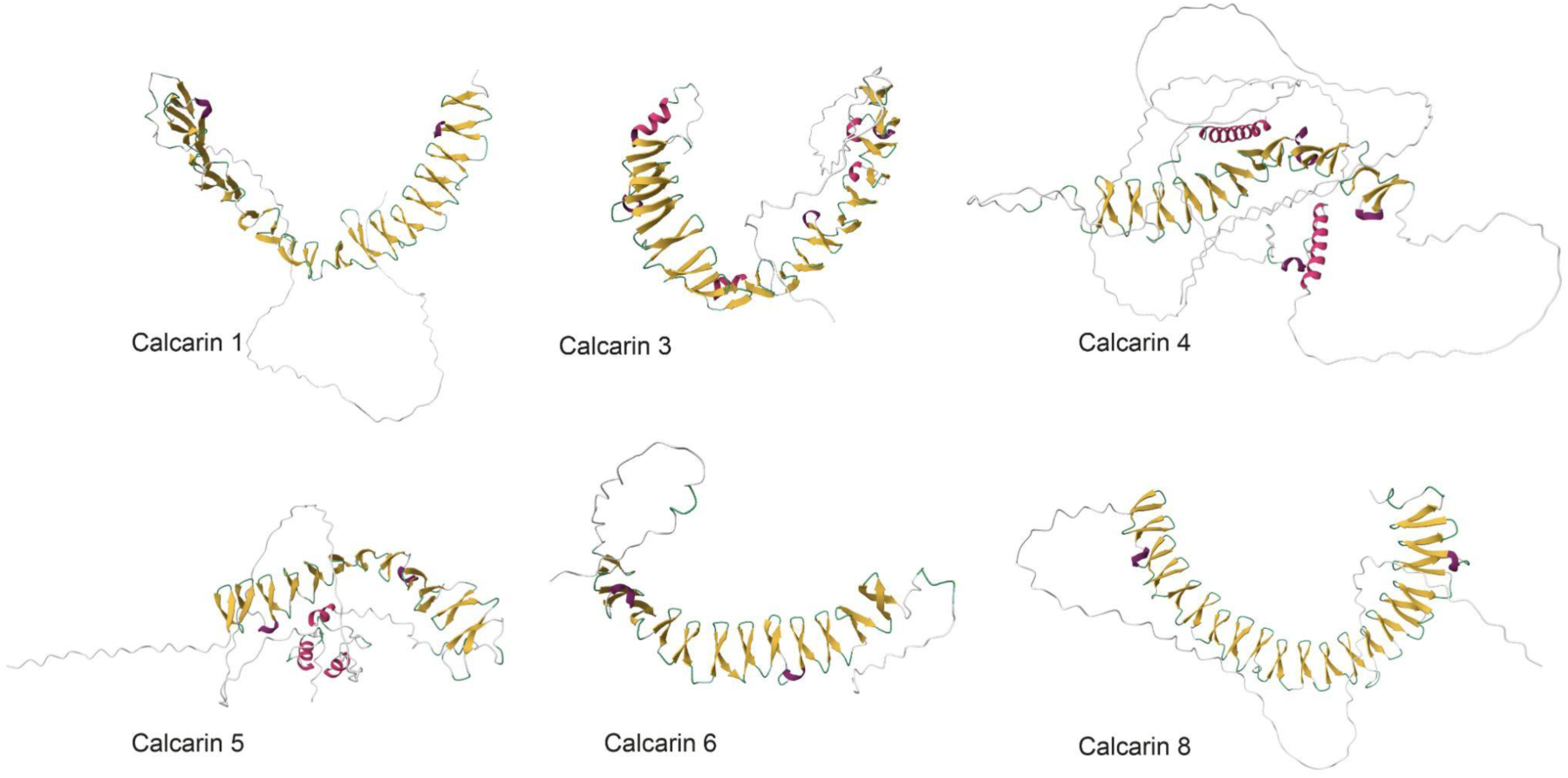
Alpha-Fold structure predictions of additional selected *Sycon ciliatum* calcarins.

**Fig. S2.**
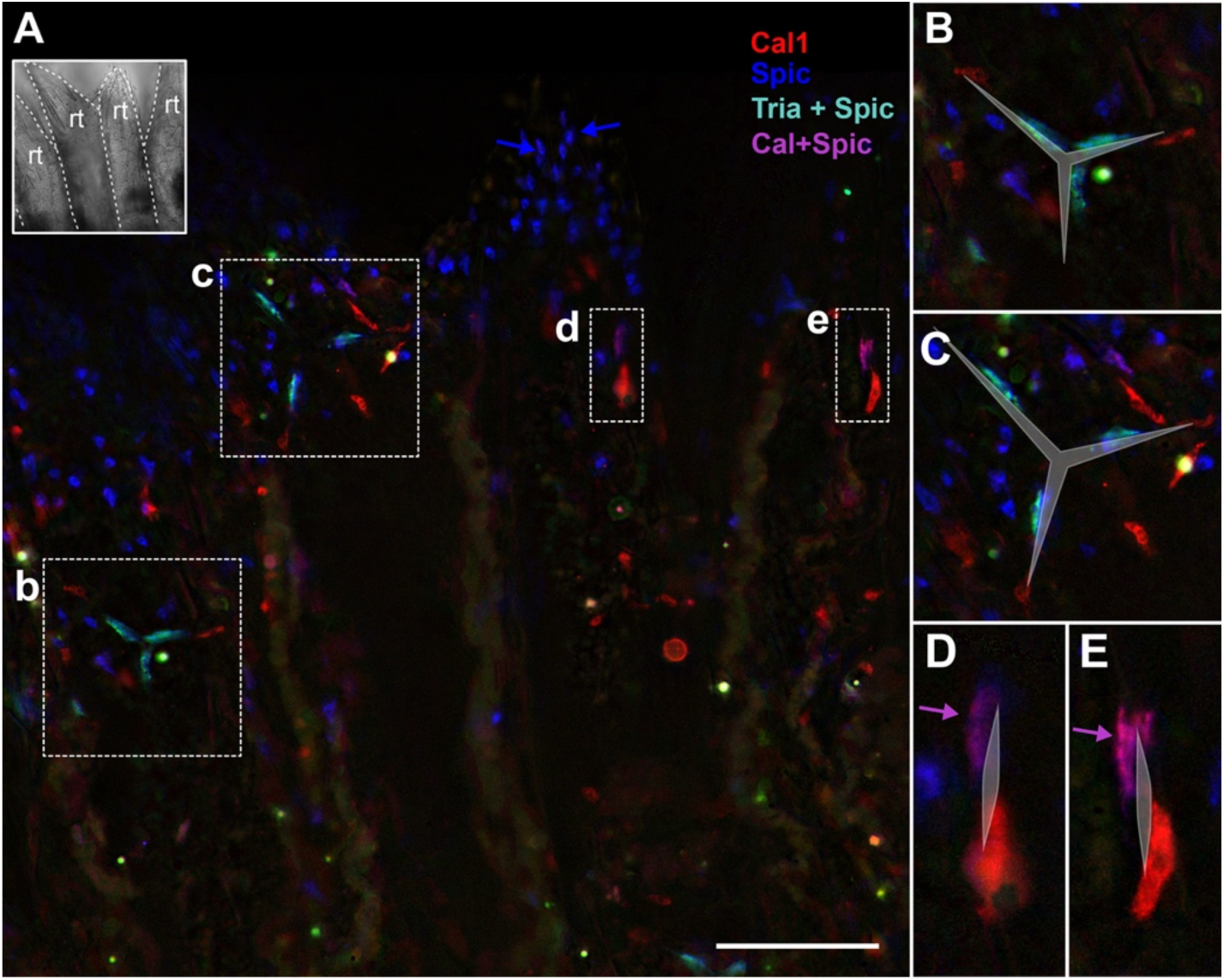
Expression of biomineralization genes in radial tubes (rt). A: Overview of fluorescent signals in radial tubes. Inset in the top left shows brightfield view for orientation. Doted boxes indicate the position of details in B-E. B-E: Details with superimposed sketches to show the original position of dissolved spicules. Triactinin and Spiculin are co-expressed in thickener cells of triactines (B & C). Spiculin additionally occurs in thickener cells of diactines at the distal end of radial tubes (e.g., arrows in A). Cal1 is expressed in founder cells of both spicule types (B-E). Co-expression of Cal1 and Spiculin marks the transition stage from founder to thickener cell (arrows in D & E). Many Spiculin signals at the distal end of radial tubes are not associated with a Cal1 signal, suggesting that Cal1 expression in diactine founder cells ceases before spicule formation is complete (A). In contrast, Cal1 signal is detected in founder cells of late triactine stages (C). Cal1= Calcarin1, Tria= Triactinin, Spic= Spiculin, rt= radial tube. Scale bar 100 µm. Images processed with LVCC.

**Fig. S3.**
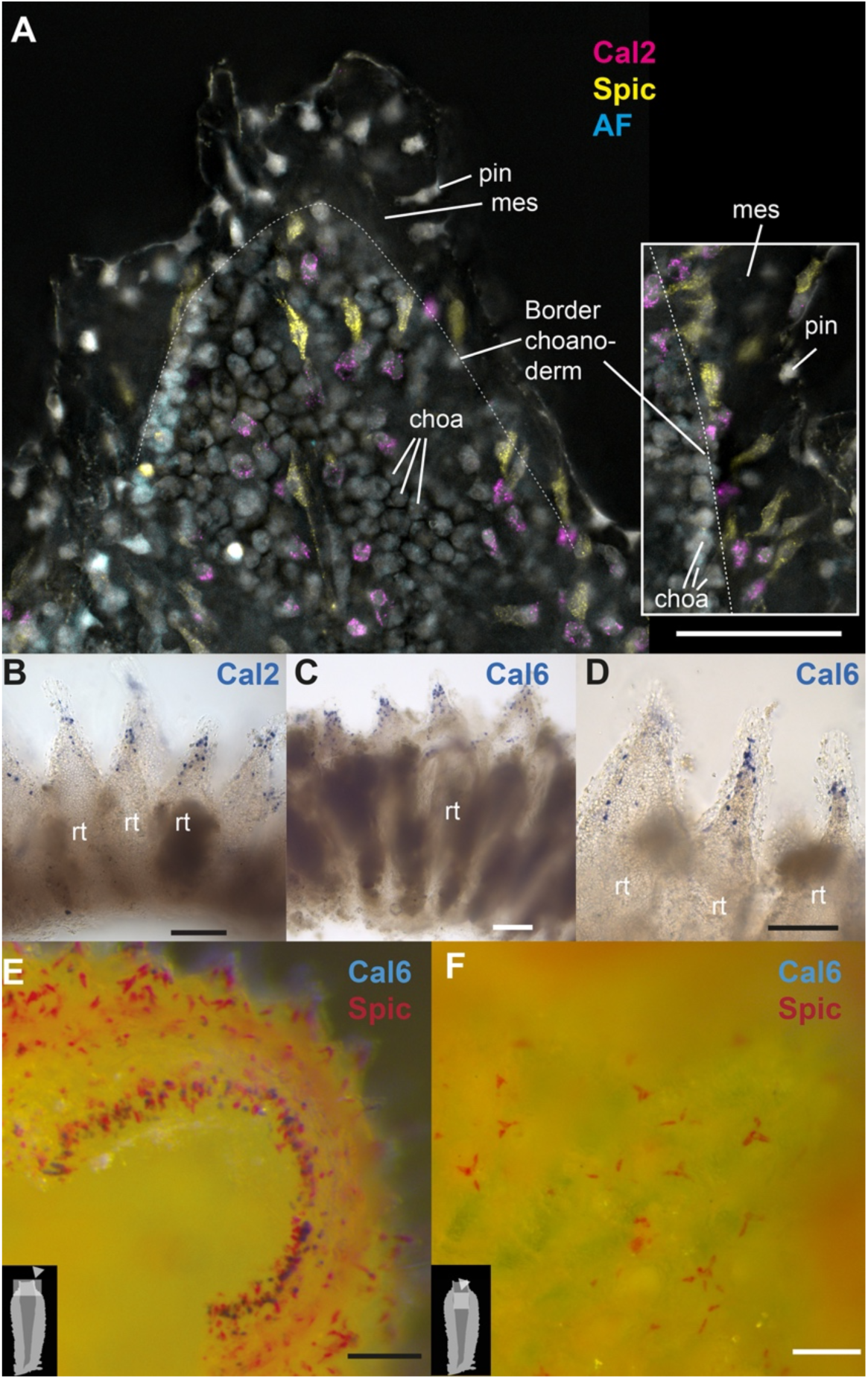
A: The radial tube’s distal end shows Cal2 expression in spherical sclerocytes closely associated with the choanoderm. Inset: View that shows the position of Cal2-expressing cells at the basal surface of the choanoderm inside the mesohyl (Images processed with LVCC, channel colors changed to cyan/magenta/yellow as described in the Extended Methods). B-D: Similar expression patterns of Cal2 and Cal6 in distal radial tubes. E: Expression of Cal6 and Spiculin around the oscular opening, F: atrial wall (no diactines) of the same individual lacks Cal6-expressing cells associated with triactine and tetractine thickener cells. AF: Autofluorescence detected with the Leica TXR filter (approx. 590–650 nm), included to help distinguish true signal from background autofluorescence observed in the FITC channel (used for Spiculin detection). **Cal:** Calcarin, **choa:** choanocytes, **mes:** mesohyl, **pin:** pinacocyte, **rt**= radial tube, **Spic:** Spiculin. Scalebar 50 µm (A), 20 µm (B-D), 100 µm (E-F).

**Fig. S4.**
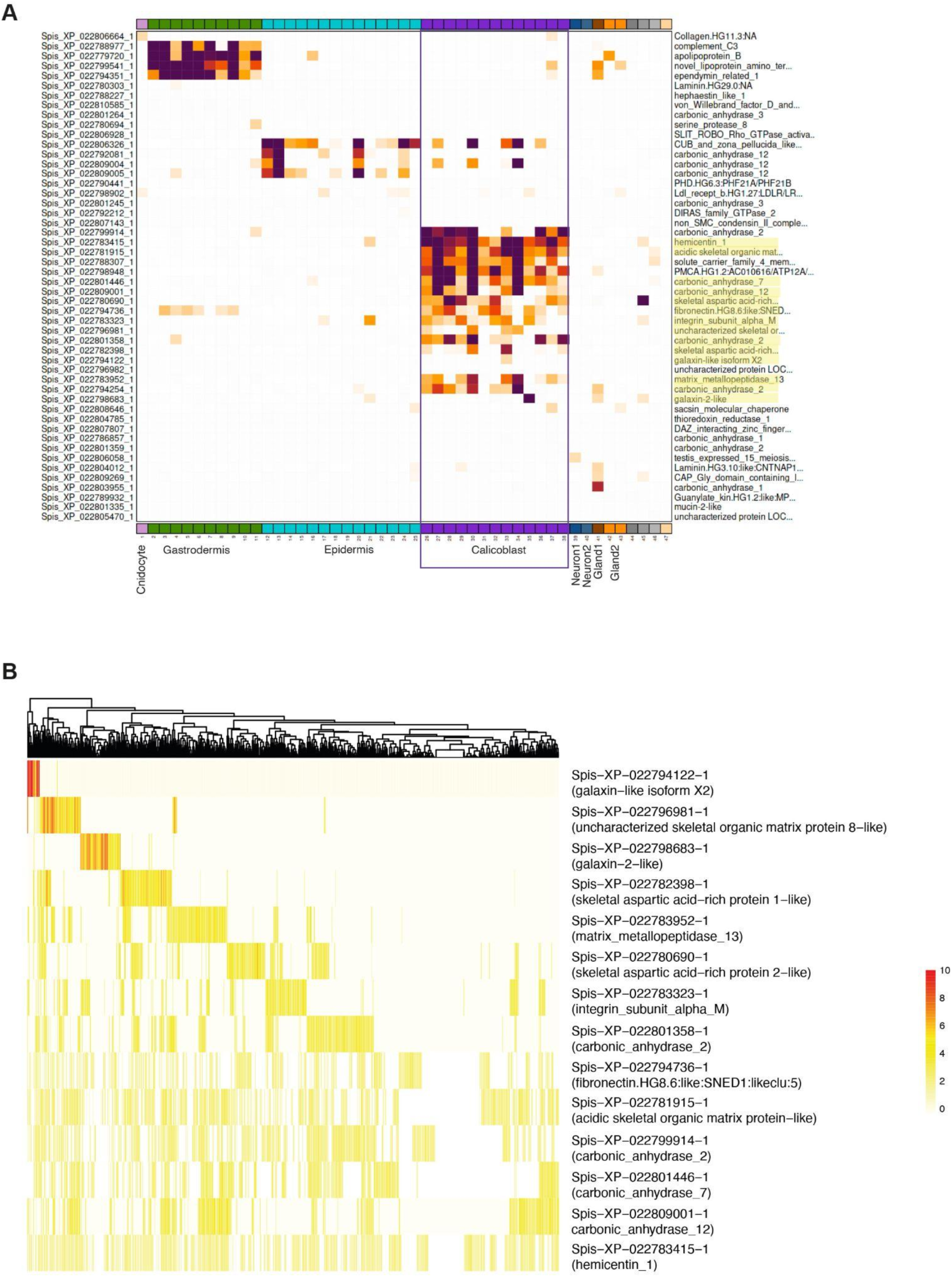
Expression of SOM proteins in cells of young *Stylophora pistillata* polyps. A. Fourteen of the known SOM proteins (Peled et al., 2020) are specifically overexpressed in calicoblast metacells. Graph obtained from https://sebe-lab.shinyapps.io/Stylophora_cell_atlas/. B. Normalized and scaled expression of 980 calicoblast cells show that several secreted SOM proteins are exclusively expressed by different calicoblast cells, suggesting a spatio-temporal expression regulation as observed in calcareous sponges.

**Fig. S5.**
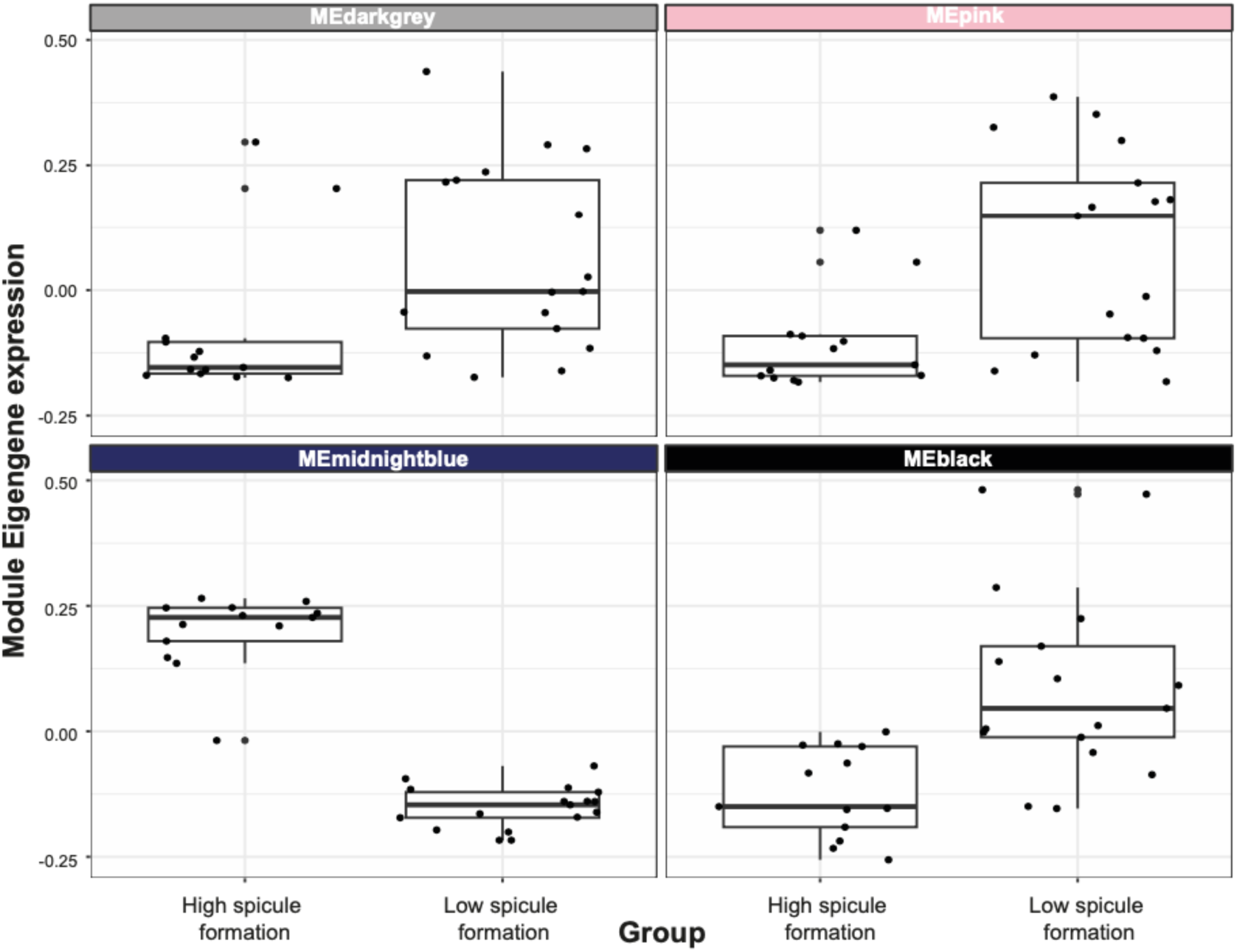
Changes of Module Eigengene expression between low spicule formation transcriptomes and high spicule formation transcriptomes. Most known biomineralization effector genes occur in MEmidnightblue.

**Fig. S6.**
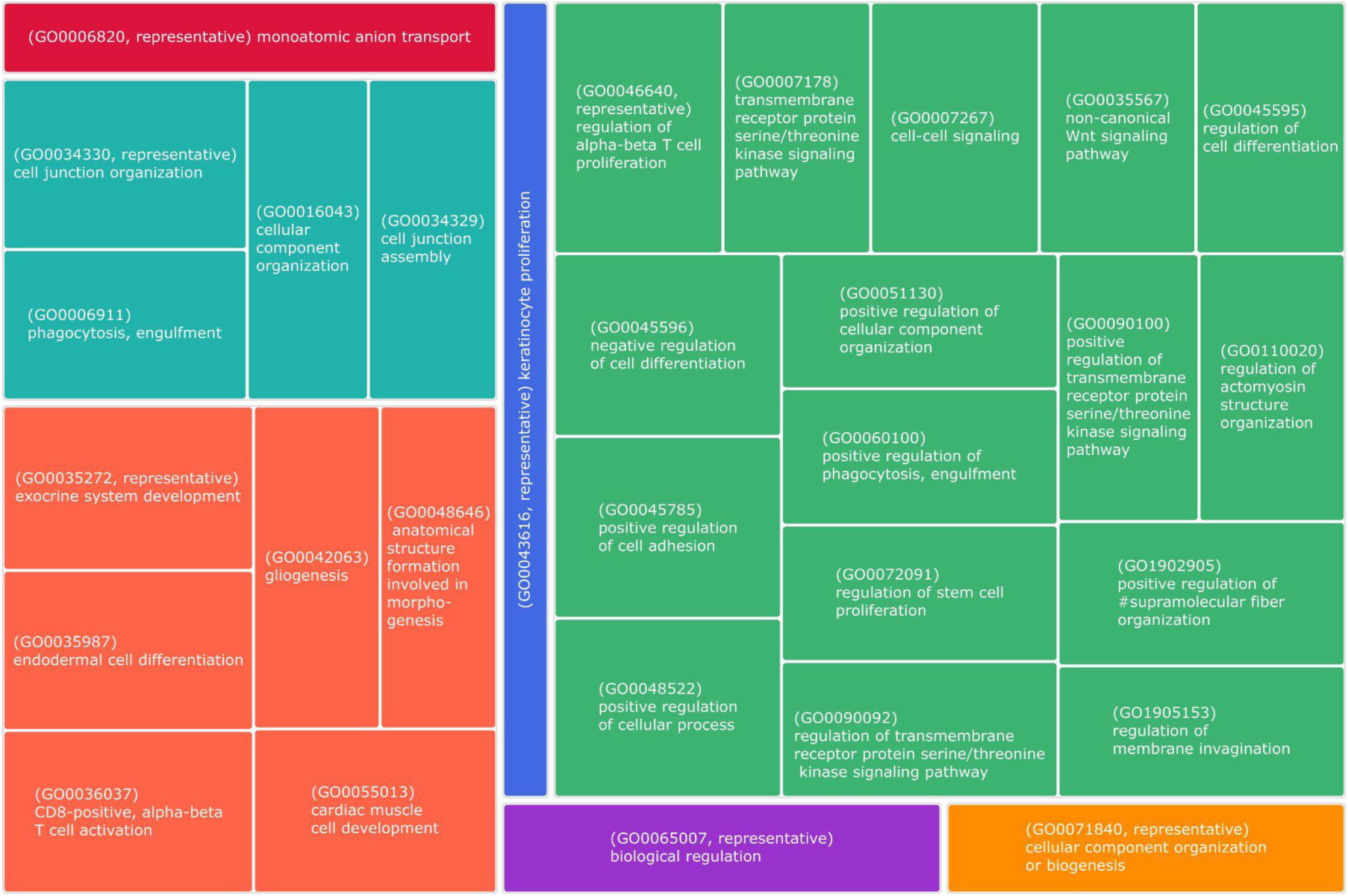
Enriched biological process GO-terms (Treemap from Revigo) of genes in meta module MEmidnightblue.

**Fig. S7.**
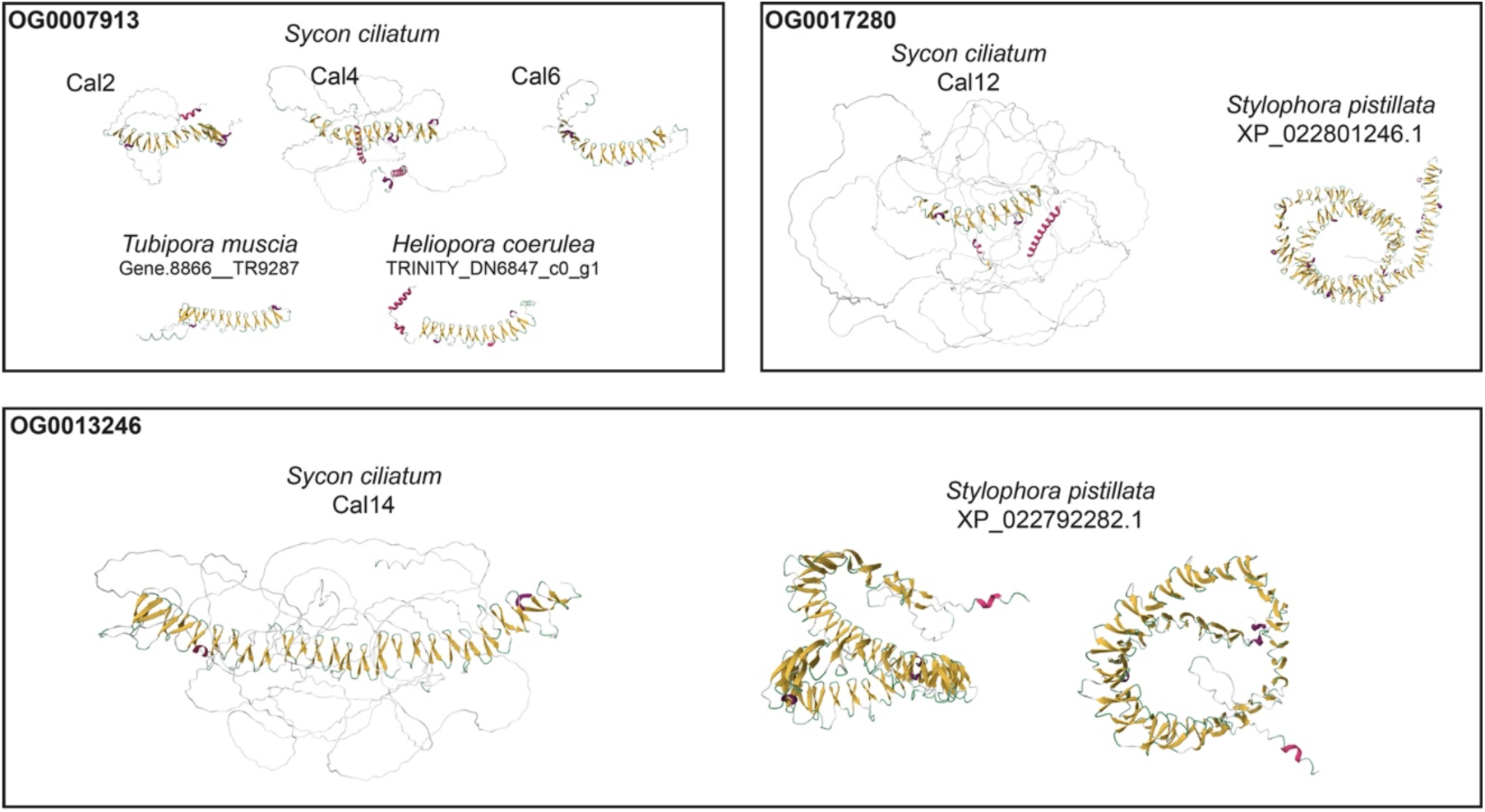
AlphaFold predictions of galaxin-like proteins from octocorals and stony corals that fall in orthogroups with ***Sycon ciliatum*** *calcarins.*.Two octocoral galaxin-like proteins exhibit the same number of beta-hairpins as ***Sycon ciliatum*** calcarins Cal2, Cal4, and Cal6 (top left). In contrast, stony coral galaxin-like proteins belonging to the same orthogroups as ***Sycon ciliatum*** Cal12 or Cal14 display considerably more beta-hairpins than their calcareous sponge counterparts (top right, bottom). Notably, the galaxin-like proteins shown here were not detected in the skeletal matrix proteomes of the respective cnidarian species.

**Fig. S8.**
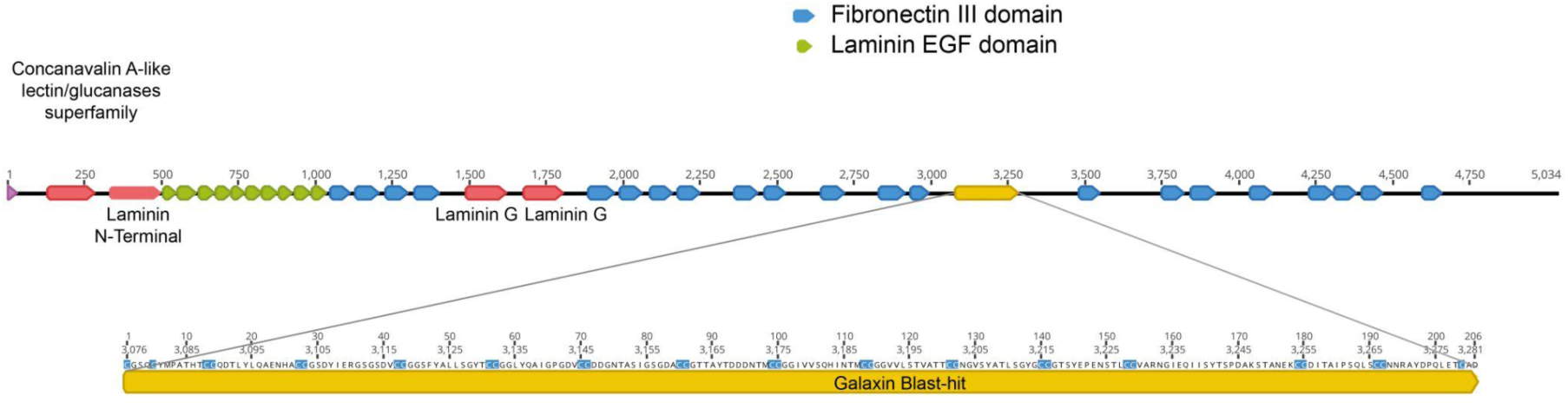
Domain structure of the only glass sponge (*Vazella pourtalesii*) protein with a blast hit for a Galaxin query (Dataset S6).

**Fig. S9.**
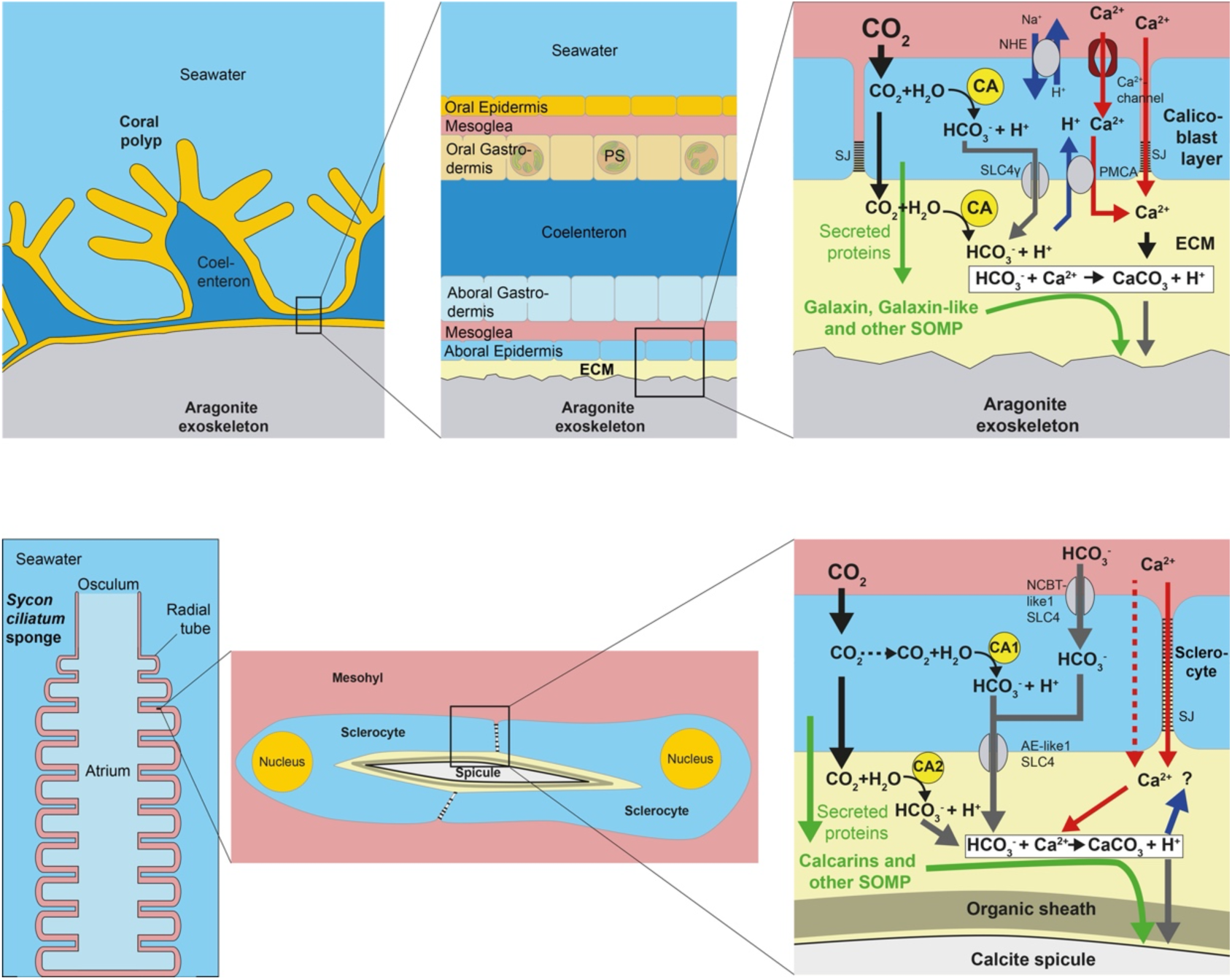
Schematic overview of skeleton formation in stony corals (top) and calcareous sponges (bottom). In stony corals, the skeleton is an extracellular exoskeleton deposited beneath the calicoblastic cell layer (the aboral epidermis), within a semi-isolated compartment known as the extracellular calcifying medium (ECM). Carbonic anhydrases in the cytosol and ECM catalyze the conversion of CO₂ to HCO₃⁻, supplying inorganic carbon for calcification. The AE-like SLC4γ transporter, localized to the apical membrane of calicoblastic cells, exports HCO₃⁻ into the ECM. Ca²⁺ reaches the ECM via both paracellular and transcellular pathways. In the paracellular route, Ca²⁺ diffuses through septate junctions between calicoblastic cells. The transcellular route involves Ca²⁺ influx channelson the basolateral membrane and plasma membrane Ca²⁺-ATPases (PMCAs) on the apical membrane, which export Ca²⁺ into the ECM while simultaneously importing protons (H⁺). To maintain intracellular pH, H⁺ is extruded from calicoblastic cells via Na⁺/H⁺ exchangers (NHEs).In the ECM, Ca²⁺ and HCO₃⁻ react to form calcium carbonate (CaCO₃), the mineral phase of the skeleton. Calicoblastic cells also secrete skeletal organic matrix proteins (SOMPs), such as galaxin and galaxin-like proteins, into the ECM, where they likely modulate crystal nucleation and growth. In calcareous sponges, the skeleton consists of calcite spicules formed by sclerocytes located in the mesohyl. Each spicule develops within an extracellular calcifying space enclosed by at least two sclerocytes (e.g., in diactine formation), which are connected by septate junctions (SJ) that seal the compartment. Inside this space, the growing spicule is surrounded by an organic sheath. Carbonic anhydrases, including mitochondrial (e.g., *Sycon ciliatum* CA1) or cytosolic forms, catalyze the conversion of CO₂ to HCO₃⁻, providing inorganic carbon for calcification. Two sclerocyte-specific SLC4 family HCO₃⁻ transporters, AE-like1 and NCBT-like1 mediate HCO₃⁻ export into and import from the calcifying space, respectively (note that their apical and basolateral localization as depicted here is speculative). Ca²⁺ is thought to enter the calcifying space via the paracellular route through junctional spaces between sclerocytes. Components of a transcellular Ca²⁺ pathway have not yet been characterized. Skeletal organic matrix proteins (SOMPs), such as calcarins, are secreted into the calcifying space, where they likely influence the biomineralization and get incorporated into the calcite spicule.

**Fig. S10.**
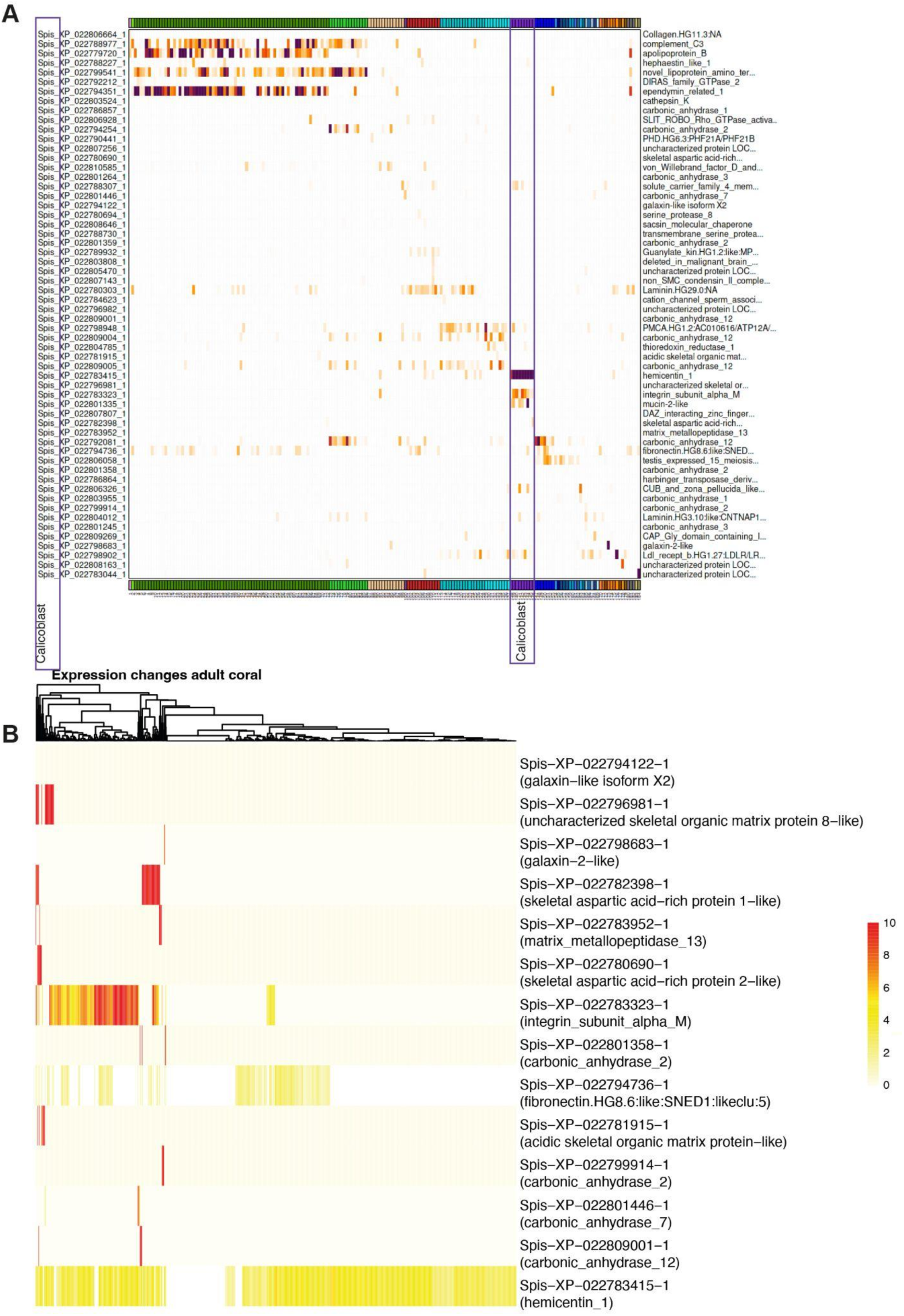
Expression of SOM proteins in adult *Stylophora pistillata* corals. A. Most calicoblast metacells did not express known SOM proteins (Peled et al., 2020).Graph obtained from https://sebe-lab.shinyapps.io/Stylophora_cell_atlas/. B. Normalized and scaled expression of the 14 SOM proteins specific to polyp calicoblasts (Fig. S4) in 896 calicoblast cells of adult corals.

**Fig. S11.**
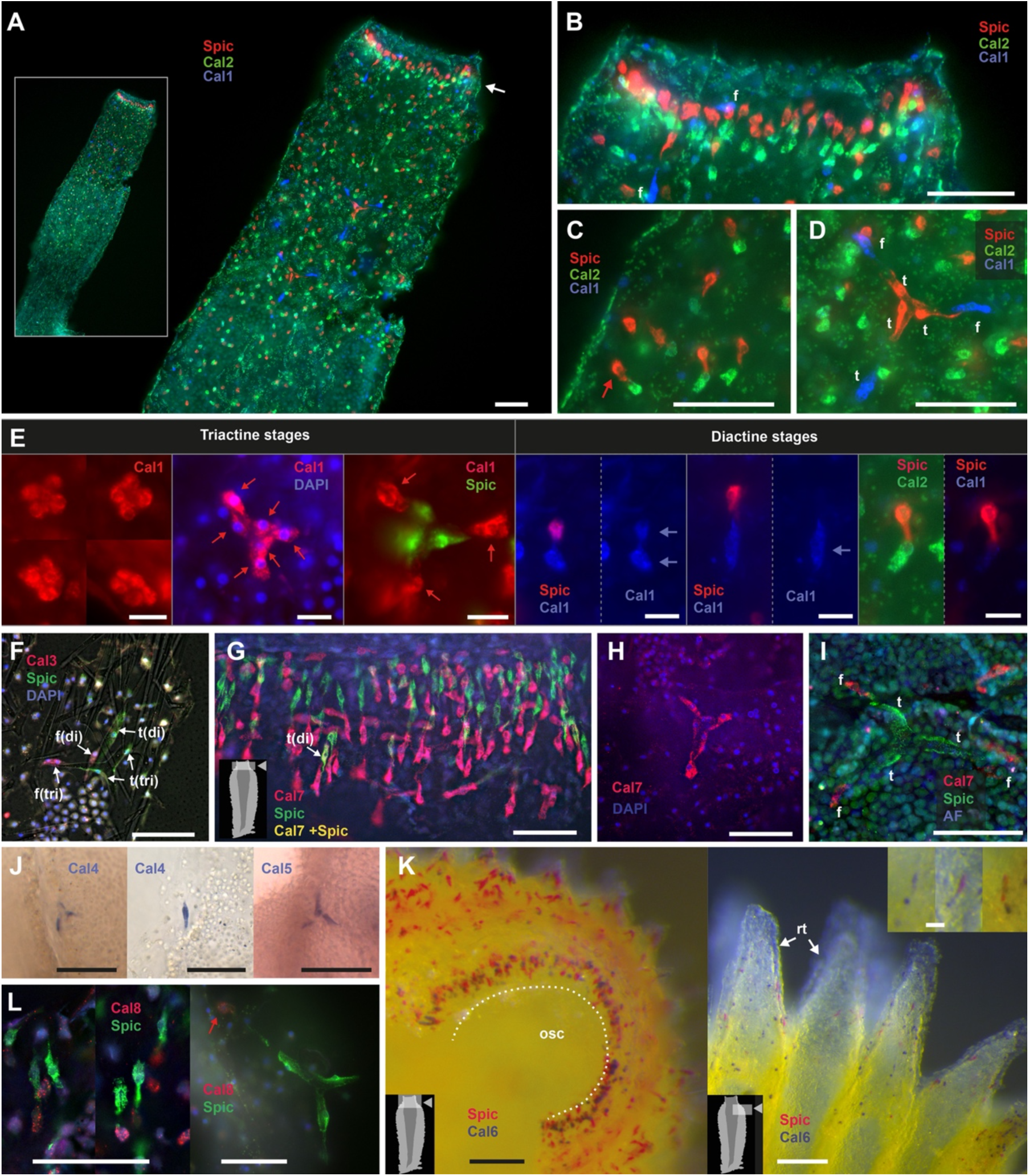
Version of Fig. 3 with the original RGB channels of the fluorescent images (A-I, L).

**Table S1.**
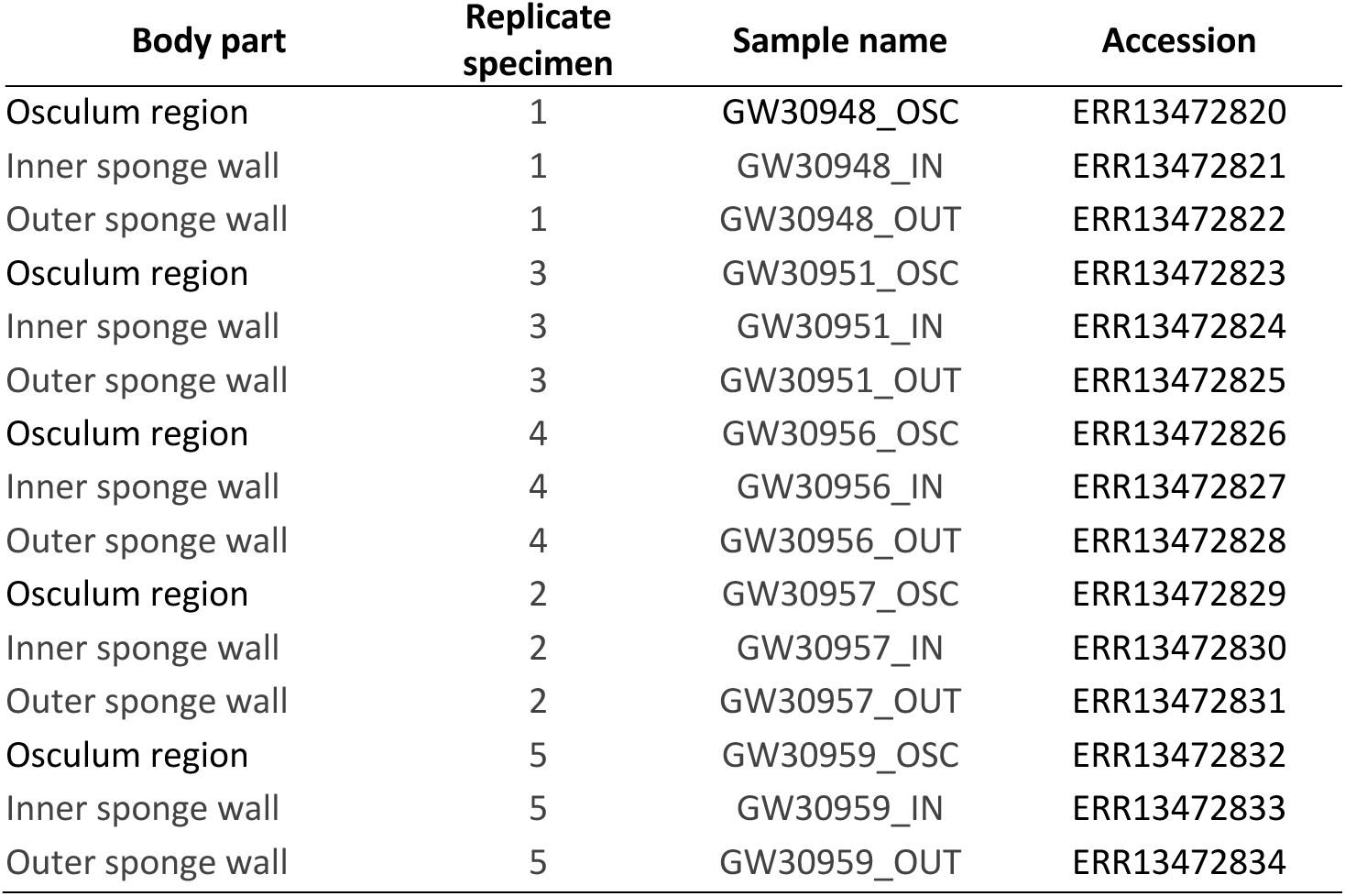
Accession numbers or RNAseq data generated for the body part dataset (BioProject PRJEB78728).

| Body part | Replicate specimen | Sample name | Accession |
| --- | --- | --- | --- |
| Osculum region | 1 | GW30948_OSC | ERR13472820 |
| Inner sponge wall | 1 | GW30948_IN | ERR13472821 |
| Outer sponge wall | 1 | GW30948_OUT | ERR13472822 |
| Osculum region | 3 | GW30951_OSC | ERR13472823 |
| Inner sponge wall | 3 | GW30951_IN | ERR13472824 |
| Outer sponge wall | 3 | GW30951_OUT | ERR13472825 |
| Osculum region | 4 | GW30956_OSC | ERR13472826 |
| Inner sponge wall | 4 | GW30956_IN | ERR13472827 |
| Outer sponge wall | 4 | GW30956_OUT | ERR13472828 |
| Osculum region | 2 | GW30957_OSC | ERR13472829 |
| Inner sponge wall | 2 | GW30957_IN | ERR13472830 |
| Outer sponge wall | 2 | GW30957_OUT | ERR13472831 |
| Osculum region | 5 | GW30959_OSC | ERR13472832 |
| Inner sponge wall | 5 | GW30959_IN | ERR13472833 |
| Outer sponge wall | 5 | GW30959_OUT | ERR13472834 |

**Table S2.** Accession for RNAseq data of regeneration experiment from Soubigou et al. (2020). Set I and set II are two regeneration experiments followed for 24 days. Additional experiments of the study were not used, because they did not include the first spicule-free stages.

| **Experiment, time** | **Accession** | **Regeneration stage** |
| --- | --- | --- |
| Set I, day 1 | SRR11617503 | No spicules |
| Set II, day 1 | SRR11617504 | No spicules |
| Set I, day 2 | SRR11617505 | No spicules |
| Set II, day 2 | SRR11617506 | No spicules |
| Set I, day 3-4 | SRR11617507 | Primmorphs with spicules (diactines) |
| Set II, day 3-4 | SRR11617508 | Primmorphs with spicules (diactines) |
| Set I, day 6-8 | SRR11617513 | Ciliated chambers |
| Set II, day 6-8 | SRR11617514 | Ciliated chambers |
| Set I, day 10-14 | SRR11617519 | Choanoderm, expanding spongocoel, pinacoderm |
| Set II, day 10-14 | SRR11617520 | Choanoderm, expanding spongocoel, pinacoderm |
| Set I, day 16-18 | SRR11617523 | Osculum opens, porocytes form ostia |
| Set II, day 16-18 | SRR11617524 | Osculum opens, porocytes form ostia |
| Set I, day 21 -24 | SRR11617526 | Juvenile |
| Set II, day 21 -24 | SRR11617527 | Juvenile |
| dissociated cells, day 0 | SRR11617528 | Adult |

**Table S3.** Source of the data used in the OrthoFinder analysis.

| Species | Accession (run, sample, study, genome) or other source | Protein predictions from |
| --- | --- | --- |
| Porifera |  |  |
| Calcarea |  |  |
| Calacronea |  |  |
| *Grantia compressa* | SRR3417193 | transcriptome* |
| *Leuconia nivea* | SRR3417190 | transcriptome* |
| *Leucosolenia complicata* | http://compagen.unit.oist.jp/datasets.html | transcriptome* |
| *Sycon ciliatum* | GCA_964019385.1 | genome** |
| *Sycon ciliatum* (Bergen) | http://compagen.unit.oist.jp/datasets.html | genome |
| *Sycon ciliatum* (Helgoland) | ERP163002 (this study) | transcriptome* |
| Calcinea |  |  |
| *Clathrina coriacea* | SRR3417192 | transcriptome* |
| *Janusya* sp. | ERR5279461 | transcriptome* |
| *Leucetta chagosensis* | SRS8111786 | transcriptome* |
| Homocleromorpha |  |  |
| *Oscarella carmela* | http://compagen.unit.oist.jp/datasets.html | genome |
| Demospongiae |  |  |
| *Ephydatia muelleri* | https://bitbucket.org/EphydatiaGenome/ephydatiagenome/downloads/ | genome |
| *Tethya wilhelma* | https://bitbucket.org/molpalmuc/tethya_wilhelma-genome/ | genome |
| *Vaceletia* sp. | SRR4423080 | transcriptome* |
| Hexactinellida |  |  |
| *Acanthascus vastus* | https://github.com/PalMuc/Aphrocallistes_vastus_genome/ | genome |
| *Vazella pourtalesii* | https://doi.org/10.6084/m9.figshare.23799351 |  |
| Cnidaria |  |  |
| Hexacorallia |  |  |
| *Acopora millepora* | GCF_013753865.1 | genome |
| *Acropora digitata* | GCF_000222465.1 | genome |
| *Stylophora pistillata* | GCF_002571385.2 | genome |
| Octocorallia |  |  |
| *Heliopora coerulea* | ERP120267 | transcriptome* |
| *Pinnigorgia flava* | ERP122203 | transcriptome* |
| *Tubipora musica* | ERR3026435 | transcriptome* |

\*re-assembled with TransPi; \*\* new gene prediction using BRAKER 3

**Table S4.**
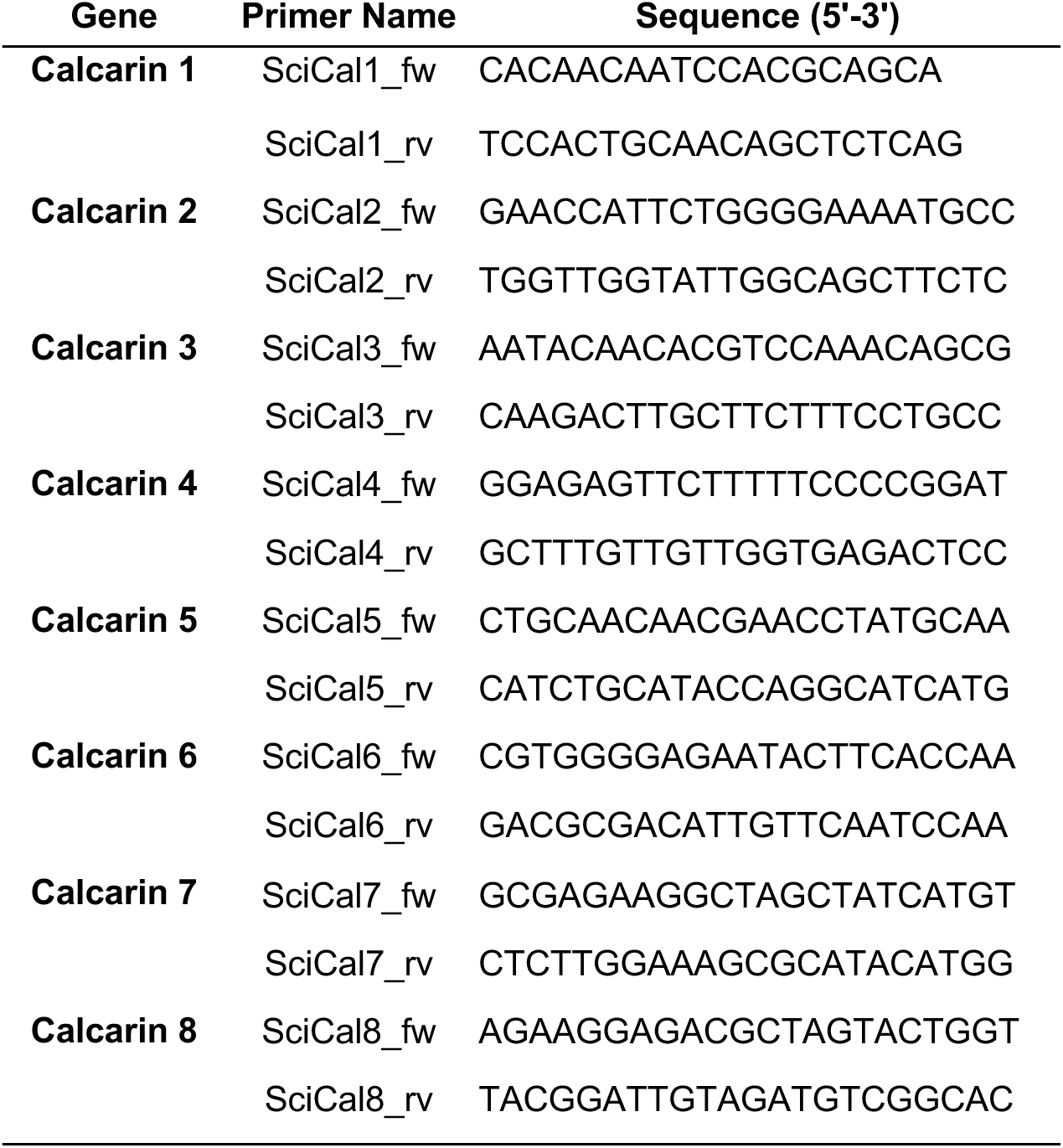
Gene-specific primer sequences for generating ISH probes

**Table S5.** HCR FISH probe sets, each consisting of 20 pairs of gene-specific probes with specific split HCR initiators. Visualization of co-expressed genes requires each target probe set to have a different HCR initiator.

| Target gene | HCR initiator |
| --- | --- |
| Triactinin | B1 |
| Spiculin | B3 |
| Calcarin 1 | B2 |
| Calcarin 2 | B1 |
| Calcarin 3 | B1 |
| Calcarin 7 | B1 |
| Calcarin 8 | B1 |
| *S.ciliatum* carbonic anhydrase 1 | B1 |

